# Evolutionary genomics reveals variation in structure and genetic content implicated in virulence and lifestyle in the genus *Gaeumannomyces*

**DOI:** 10.1101/2024.02.15.580261

**Authors:** Rowena Hill, Michelle Grey, Mariano Olivera Fedi, Daniel Smith, Sabrina J. Ward, Gail Canning, Naomi Irish, Jade Smith, Vanessa E. McMillan, Jess Hammond, Sarah-Jane Osborne, Tania Chancellor, David Swarbreck, Neil Hall, Javier Palma-Guerrero, Kim E. Hammond-Kosack, Mark McMullan

## Abstract

*Gaeumannomyces tritici* is responsible for take-all disease, one of the most important wheat root threats worldwide. High-quality annotated genome resources are sorely lacking for this pathogen, as well as for the closely related antagonist and potential wheat take-all biocontrol agent, *G. hyphopodioides*. As such, we know very little about the genetic basis of the interactions in this host-pathogen-antagonist system. Using PacBio HiFi sequencing technology we have generated nine near-complete assemblies, including two different virulence lineages for *G. tritici* and the first assemblies for *G. hyphopodioides* and *G. avenae* (oat take-all). Genomic signatures support the presence of two distinct virulence lineages in *G. tritici* (types A and B), with A strains potentially employing a mechanism to prevent gene copy-number expansions. The CAZyme repertoire was highly conserved across *Gaeumannomyces*, while candidate secreted effector proteins and biosynthetic gene clusters showed more variability and may distinguish pathogenic and non-pathogenic lineages. A transition from self-sterility (heterothallism) to self-fertility (homothallism) may also be a key innovation implicated in lifestyle. We did not find evidence for transposable element and effector gene compartmentalisation in the genus, however the presence of *Starship* giant transposable elements likely contributes to genomic plasticity in the genus. Our results depict *Gaeumannomyces* as an ideal system to explore interactions within the rhizosphere, the nuances of intraspecific virulence, interspecific antagonism, and fungal lifestyle evolution. The foundational genomic resources provided here will enable the development of diagnostics and surveillance of understudied but agriculturally important fungal pathogens.

## INTRODUCTION

*Gaeumannomyces* is a broadly distributed genus of *Poaceae* grass-associated root-fungi (Hernández-Restrepo et al. 2016), best known for the species *Gaeumannomyces tritici* (*Gt*) which causes take-all disease, the most serious root disease of wheat (Palma-Guerrero et al. 2021). *Gaeumannomyces* is a comparatively understudied genus despite belonging to the *Magnaporthales*, an economically important order of pathogens including the rice and wheat blast fungus *Pyricularia oryzae* (syn. *Magnaporthe oryzae* (Zhang et al. 2016)). This is perhaps due to a historical research bias towards above-ground pathogens, in part simply due to the fact that characteristic symptoms of root pathogen diseases are hidden from view (Raaijmakers et al. 2009; Balmer and Mauch-Mani 2013). Recently the rhizosphere has received more research attention as its key role in plant health and productivity has become apparent (van der Heijden et al. 2008). There have also been considerable difficulties in producing a reliable transformation system for *Gt*, preventing gene disruption experiments to elucidate function (Freeman and Ward 2004).

Although genetic studies of *Gt* have been limited, single-locus phylogenetic analyses of *Gt* have consistently recovered two distinct lineages within the species (Daval et al. 2010), which we will refer to using the ‘A/B’ characterisation established by Freeman et al. (2005) based on *ITS2* polymorphism. Although very little is known about the dynamics of these two lineages, each is found across the world and both lineages persistently co-occur in the same field, prompting the suggestion that the two lineages may actually be cryptic species (Daval et al. 2010; Palma-Guerrero et al. 2021). Although variation within lineages is high, there is also some evidence that type A strains are more virulent (Bateman et al. 1997; Lebreton et al. 2004, 2007), which is a major impetus for improving our understanding of these two lineages. The sister species to *Gt*, *G. avenae* (*Ga*), can also infect wheat, but is not the predominant agent of wheat take-all, and is distinguished by the fact that production of avenacinase enables *Ga* to additionally infect oat roots (Osbourn et al. 1991; Bowyer et al. 1995).

*Magnaporthales* are also home to several commensal and/or mutualistic fungi (Xu et al. 2014), including those with the potential to inhibit take-all (Chancellor 2022). For instance, *G. hyphopodioides* (*Gh*) — a species closely related to *Gt* that also grows on wheat roots— is not only non-pathogenic, but actually capable of suppressing take-all to varying degrees (Osborne et al. 2018). It is now apparent that prior *Gh* colonisation primes the host plant’s immune response (Chancellor et al. 2023), a mechanism that has been reported in various other plant–microbe interactions associated with disease prevention (Van Wees et al. 2008; Zamioudis and Pieterse 2012). This has prompted interest in *Gh* as a potential biocontrol agent, for instance by adding *Gh* inoculant to wheat seedstock via seed coating (Accinelli et al. 2016) and/or selecting for wheat cultivars that support enhanced levels of *Gh* root system colonisation (Osborne et al. 2018). Novel disease prevention approaches for take-all are especially desirable as up to 30% of *Gt* strains are found to be naturally resistant to the seed-dressing fungicide routinely used to treat take-all, silthiofam (Freeman et al. 2005).

Understanding the genetic machinery underpinning virulence and lifestyle in *Gaeumannomyces* has previously been hampered by a lack of genomic data. Prior to the present study, a single annotated *Gt* assembly (strain R3-111a-1), sequenced using the 454 platform, was available on NCBI (accession GCF_000145635.1) (Okagaki et al. 2015) – one other more recent PacBio assembly has been released for the same strain, but remains unannotated (GCA_016080095.1). This scarcity of genomic resources has not only limited our understanding of the genetics of the system, but also accounts for a lack of molecular diagnostics for take-all. Given the increase in research activities since 2005 following the production of genomic resources for *P. oryzae* (Sperr 2023; Dean et al. 2005), we are optimistic that providing similar high-quality assemblies for *Gaeumannomyces* species will bolster research efforts in the global take-all community.

Here, we have addressed the gap in genomic resources for *Gaeumannomyces* by generating near-complete assemblies for nine strains, including both type A and B *Gt* lineages and the first assemblies for *Gh* and *Ga*. Using an evolutionary genomics approach, we identified variation in structure as well as gene features known to be involved in plant-fungal interactions — candidate secreted effector proteins (CSEPs), carbohydrate-active enzymes (CAZymes) and biosynthetic gene clusters (BGCs) — to address the questions: (1) Are there genomic signatures distinguishing *Gt* A/B virulence lineages? (2) How do gene repertoires differ between pathogenic *Gt* and non-pathogenic *Gh*? and (3) Is there evidence of genome compartmentalisation in *Gaeumannomyces*? In the process of doing so, we also identified giant cargo-carrying transposable elements belonging to the recently established *Starship* superfamily (Gluck-Thaler et al. 2022).

## RESULTS

### Evidence of greater take-all severity caused by *G. tritici* type A strains

As the five *Gt* strains sequenced in this study included representatives of both the type A and B lineages, we performed a season long inoculation experiment to determine the relative capacity for each strain to cause take-all disease symptoms. From general visual inspection, inoculation of *Gt*A strains into the highly susceptible winter wheat cultivar Hereward resulted in notably depleted roots compared to a control and, to a lesser extent, *Gt*B strains (Fig. 1a). Inoculation with *Gt*A strains also resulted in a visible reduction of overall plant size compared to the control, while *Gt*B-inoculated plants were less easily distinguished from the control (Fig. 1b). Although above- and below-ground characteristics of wheat varied depending on *Gt* strain, our statistical analysis showed that the *Gt*A strains had a greater capacity to reduce plant height and reduce root length, and both *Gt*A strains consistently produced the greatest root disease symptoms, i.e. highest Take-all Index (TAI) scores (Bateman et al. 2004) (Fig. 1c). Furthermore, five out of six wheat plants that died during the experiment were inoculated with *Gt*A strains. Several characteristics were inconsistently affected by *Gt* inoculation, including mean floral spike (ear) length; dried root biomass; number of roots; and number of roots per tiller.

**Figure 1.**
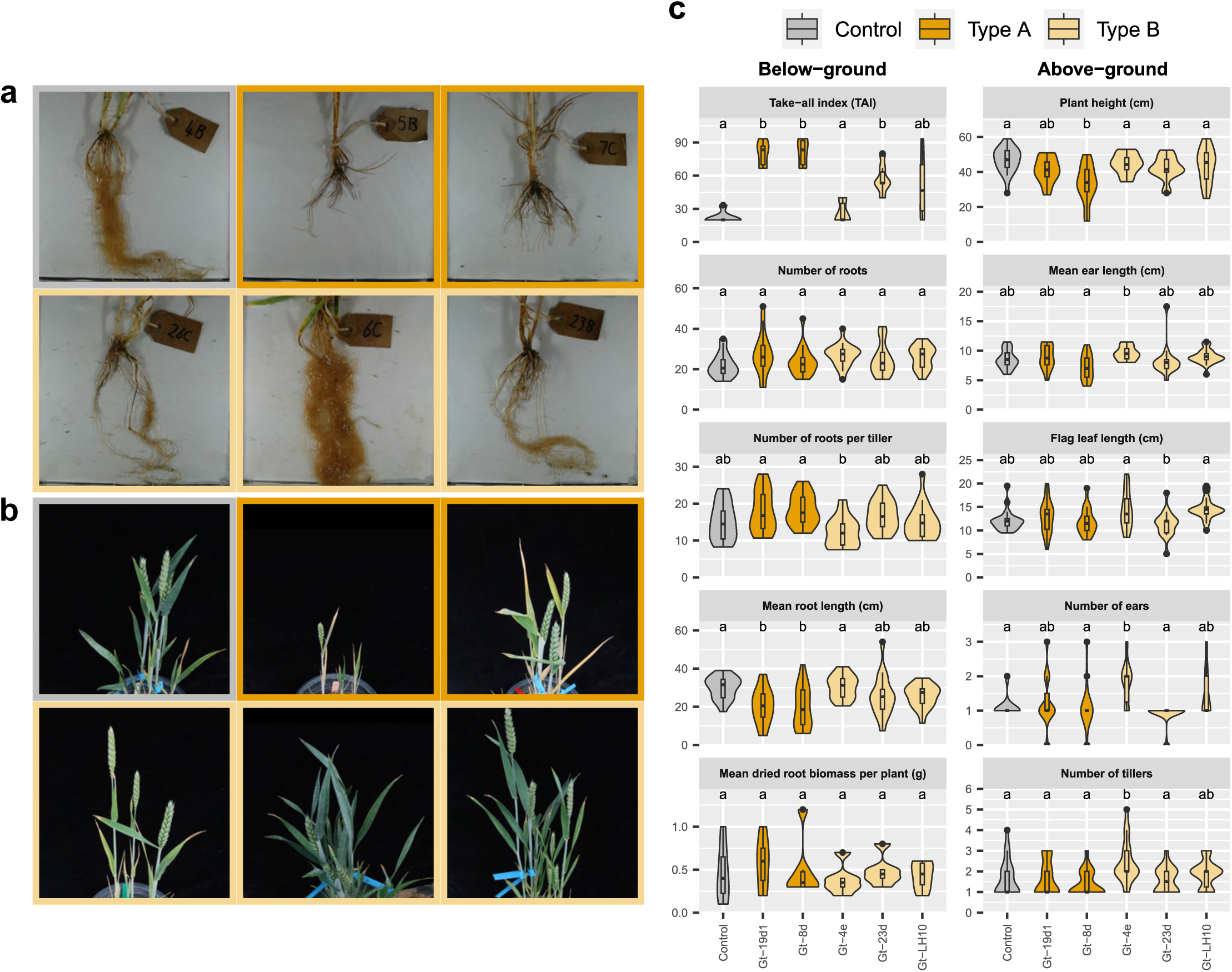
Intraspecific variation in *Gaeumannomyces tritici* (*Gt)* virulence assessed from inoculation of wheat plants. Representative photos of wheat roots (**a**) and above-ground features (**b**) following inoculation treatment. Inoculated strains from top left to bottom right: no *Gt* (control), Gt-8d, Gt-19d1, Gt-23d, Gt-4e and Gt-LH10. **c** Box and violin plots showing the impact of the five *Gt* strains sequenced in this study on above- and below-ground characteristics in winter wheat. Control, *Gt* type A and type B groups are indicated by different colours. Strains with a significant mean difference for the characteristic as calculated by either the Tukey HSD or Games-Howell test are shown by letters above the plots.

### Nine near-complete *Gaeumannomyces* assemblies, including first genome assemblies for *G. avenae* and *G. hyphopodioides*

We used PacBio HiFi sequencing technology to produce highly contiguous genome assemblies for five *Gt*, two *Gh* and two *Ga* strains. All nine assembled genomes had N50 values of more than 4 Mb (Supplemental Table S1), a 100-fold increase on the N50 of the existing annotated *Gt* RefSeq assembly (NCBI accession GCF_000145635.1). In addition, transcriptomes were sequenced for all nine strains to inform gene prediction, and between 22–29% of annotated gene models had two or more isoforms across all strains (Supplemental Fig. S1). Contigs corresponding to mitochondrial genomes were identified from all assemblies (Supplemental Table S1), however circularisation was only successfully detected for two strains (Gt-23d and Ga-CB1). For most strains the overall mitogenome size, GC content and number of genes fell within the expected range for ascomycetes (Fonseca et al. 2021), however the mitogenome assembly for Gt-LH10 is likely incomplete, as it was a third of the size of the other *Gt*B strains, and only had 23 genes annotated compared to the 38–40 genes found for all other strains (Table S1).

Combined GENESPACE (Lovell et al. 2022) and telomere prediction results suggested six chromosomes for *Gaeumannomyces* (Fig. 2), one less than *P. oryzae* (Dean et al. 2005). Telomere-to-telomere sequences were assembled for at least five out of six pseudochromosomes for most strains. By plotting GC content alongside transposable element (TE) and gene density, we also identified AT- and TE-rich but gene-poor regions, which are putative candidates for centromeres (Supplemental Fig. S2). Some of these regions additionally correspond well with points of fragmentation in other strains, presumably due to the difficulties associated with assembly of such highly repetitive regions. Other than these occasional splits into two fragments, in most cases pseudochromosomes were entire, the exception being Gh-1B17 pseudochromosome 2 which was fragmented across five contigs.

**Figure 2.**
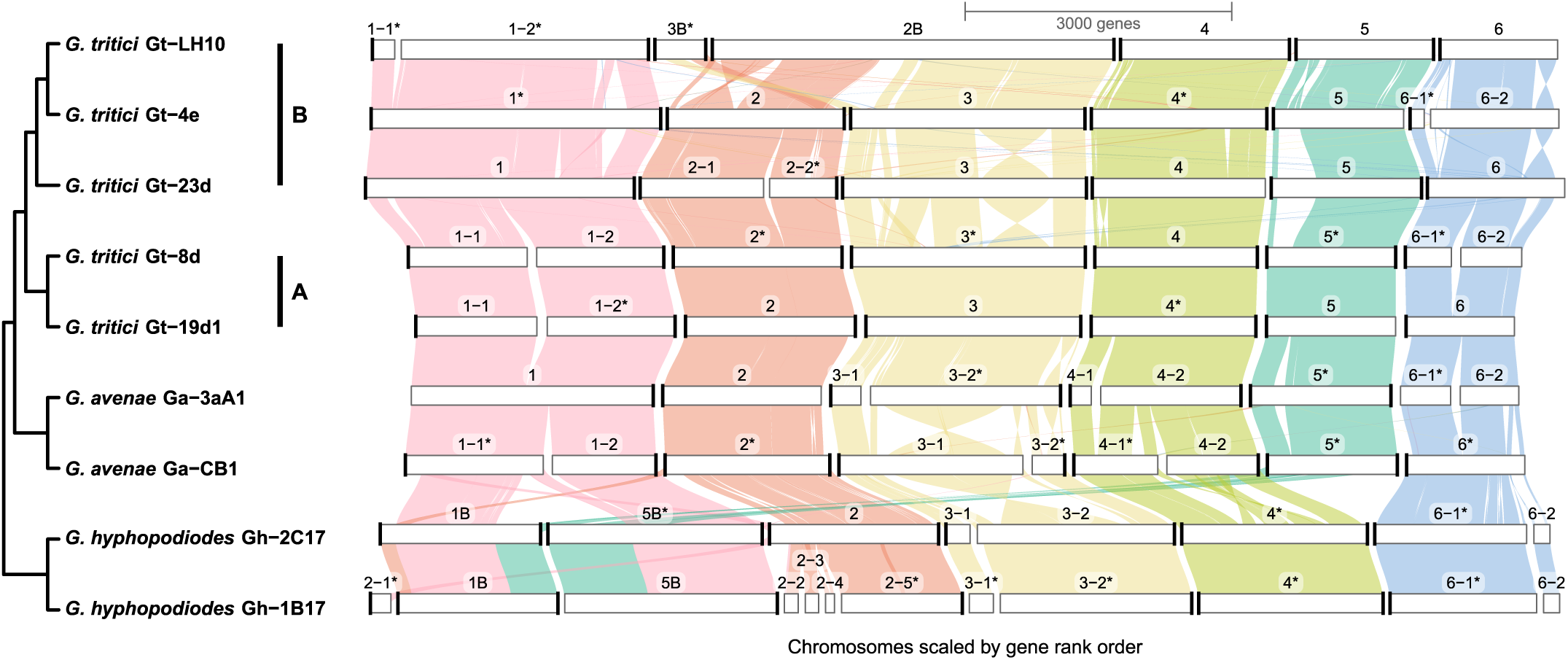
GENESPACE plot showing synteny across the nine *Gaeumannomyces* strains. A/B lineages are indicated for *G. tritici* strains. Fragments are labelled with numbers corresponding to pseudochromosomes, and an asterisk indicates that a fragment was inverted in the visualisation. Black bars on the ends of fragments indicate telomeres predicted using Tapestry.

Both *Gt*A and, to a slightly lesser degree, *Gt*B were broadly syntenic across whole pseudochromosomes, with the exception of a major chromosomal translocation between pseudochromosomes 2 and 3 in Gt-LH10 (Fig. 2). Visualisation of the spanning reads and coverage across the regions of the apparent translocation suggests the depicted arrangement is correct and not an artefact due to misassembly (Supplemental Fig. S3a), moreover there was no evidence of a block of repeats consistent with a telomere anywhere but at the ends of the pseudochromosomes (Supplemental Fig. S3c). *Ga* was also largely syntenic with *Gt*, although there were a number of inversions in Ga-CB1 pseudochromosome 3 (Fig. 2). The more distantly related *Gh* showed chromosomal translocations involving pseudochromosomes 1, 2 and 5, which were again supported by spanning reads and the absence of intrachromosomal telomeric repeats (Supplemental Fig. S3b, c).

### No evidence for significant colocalisation of transposable elements and effectors

Compartmentalisation of effectors within genomic regions enriched in transposable elements (TEs) has previously been reported for various fungal phytopathogens (Dong et al. 2015). In all the *Gaeumannomyces* strains sequenced here, however, we did not observe that predicted CSEPs were more likely to occur in regions of high TE density (Fig. 3a). We found a weak significant positive correlation between CSEP density and TE density for a minority of strains, however the scatterplot and local polynomial regression lines were unconvincing (Fig. 3b). CSEP density was more frequently found to significantly correlate with gene density, although this was still only a weak association (Fig. 3b). For all but one strain, there was no significant difference in mean distance to closest TE for CSEPs versus other genes (Fig. 3c). For strain Gt-19d1, the mean distance from a CSEP to the closest TE was marginally lower (10,036 bp) than for other genes (12,565 bp), which permutation analysis confirmed was closer than expected based on the overall gene universe (p=0.03), although this only remained significant for pseudochromosomes 2 and 6 when testing pseudochromosomes separately (Supplemental Fig. S4a). Individual pseudochromosomes for other strains also had lower than expected CSEP–TE distances, but with low z-scores (a proxy for strength) across the board. Comparing across strains, mean gene–TE distance was significantly different both within and between lineages, and lowest in *Gt*B (Fig. 3c). Within *Gt*B, Gt-LH10 had significantly lower mean gene–TE distance, and the same strain has also undergone an apparent expansion in total number of TEs compared to all other strains (Supplemental Fig. S5).

**Figure 3.**
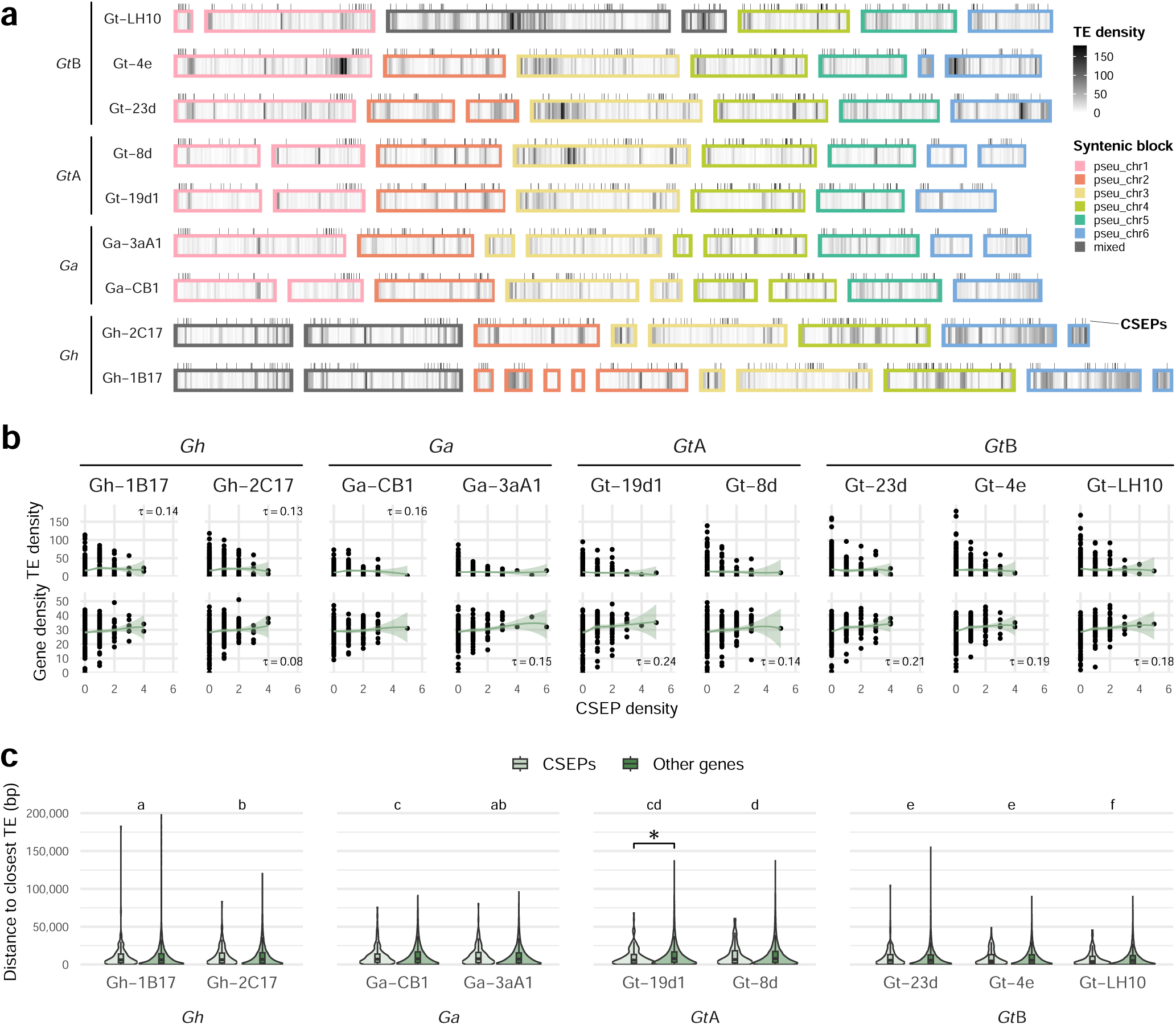
The relationship between candidate secreted effector proteins (CSEPs) and transposable elements (TEs) in *Gaeumannomyces*. **a** TE density (per 100,000 bp) and the location of CSEPs (black ticks) across fragments. Fragments are ordered syntenically according to GENESPACE (Fig. 2). **b** Scatterplot showing the relationship between CSEP density versus TE and gene density (per 100,000 bp) with local polynomial regression lines (ggplot2 function geom_smooth, method = “loess”). Significant correlation is indicated with Kendall’s tau (τ). **c** Box and violin plots showing the distance of genes to the closest TE, with CSEPs and other genes distinguished by colour. An asterisk indicates where a Wilcoxon rank sum test found the mean TE distance to be significantly different for CSEPs versus other genes. Strains with a significant mean difference in overall gene-TE distance as calculated by the Games-Howell test are shown by letters above the plots.

Although CSEPs were not broadly colocalised with TEs, we did observe that they appeared to be non-randomly distributed in some pseudochromosomes (Fig. 3a). Permutation analyses confirmed that overall CSEPs were significantly closer to telomeric regions in all strains (p=<0.008), although by testing pseudochromosomes separately we found that this pattern varied across the genome (Supplemental Fig. S4b). CSEPs on pseudochromosomes 1, 2 and 5 were consistently closer to telomeric regions, whereas for pseudochromosomes 3 and 4 CSEPs were no closer than expected based on the gene universe. CSEPs were also closer to telomeres in pseudochromosome 6, but only in *Gt* strains.

### Core gene content in *Gaeumannomyces*

The total number of genes was relatively similar for all strains, although, as indicated in Fig. 2, *Gt*B and *Gh* strains had 3–6% more genes than *Gt*A or *Ga* (Fig. 4a). *Gt*A and *Gt*B had a very similar number of CSEPs, CAZymes and BGCs, however, and more CSEPs and BGCs than either *Ga* or *Gh*. Almost all total genes, CSEPs and CAZymes were core or soft-core (i.e. present in all but one strain) in *Gt*, while there was a greater proportion of BGCs that were accessory or strain-specific. From a pangenome perspective, the core gene content for *Gt* from sampling these five strains amounted to ∼10,000 genes (Fig. 4b), which equates to ∼88% of genes per strain being core, consistent with reports in other fungi (McCarthy and Fitzpatrick 2019). The majority of BUSCO genes found to be missing in the assemblies were missing from all strains (Supplemental Fig. S6), suggesting that they are not present in the genus, rather than being missed as a result of sequencing or assembly errors. Three of these 18 missing core genes belonged to the *Snf7* family, which is involved in unconventional secretion of virulence factors in fungi (da C. Godinho et al. 2014), and is essential for pathogenicity in *P. oryzae* (Cheng et al. 2018). The next greatest set of missing BUSCOs (8) also seemed to be lineage specific – i.e. missing in *Gh* but present in *Gt*/*Ga* (Supplemental Fig. S6).

**Figure 4.**
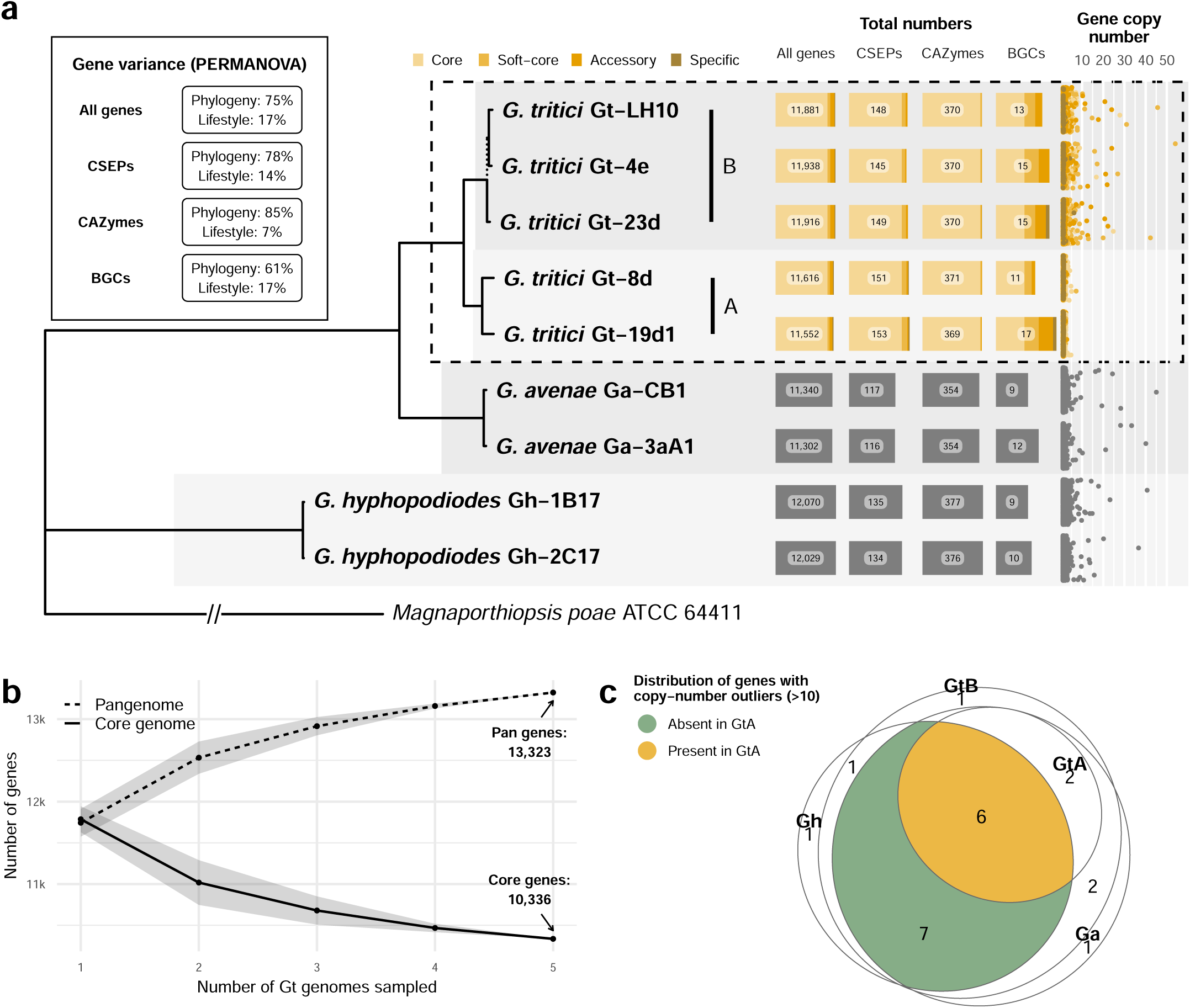
Summary of predicted gene content for the *Gaeumannomyces* strains reported in this study. **a** Number of total genes, candidate secreted effector proteins (CSEPs), carbohydrate-active enzymes (CAZymes) and biosynthetic gene clusters (BGCs) for each *Gaeumannomyces* strain. The A/B lineages are indicated for *Gaeumannomyces tritici* (*Gt*) strains. The dashed line in the phylogeny indicates bootstrap support <70 found within the *Gt*B lineage (see Supplemental Fig. S13b for the full genome-scale *Gaeumannomyces* species tree). The *Gt* pangenome (within dashed box) is categorised as core (present in all strains), soft-core (present in all but one strain), accessory (present in at least two strains) and specific (present in one strain). The lefthand inset box shows the results of PERMANOVA statistical tests which calculate the descriptive power of relatedness (phylogeny) versus lifestyle categorisation (*Gt* and *G. avenae* as pathogenic in wheat, *G. hyphopodioides* as non-pathogenic) on gene variance. Gene copy-number is shown on a scatterplot to the right, with points jittered vertically to improve visualisation. **b** Accumulation curves of pan and core genes for the *Gt* pangenome (Siozios 2021). **c** Euler diagram summarising whether high copy-number genes in each lineage are present but in low copy-number in *Gt*A, or completely absent.

The avenacinase gene required for virulence on oat roots (Osbourn et al. 1991; Bowyer et al. 1995) was identified in all strains in a conserved position on pseudochromosome 4 (Supplemental Fig. S7a). Two mating-type (MAT) loci were identified in *Gt* and *Ga*, with homologues of *Pyricularia grisea MAT1-1* and *MAT1-2* idiomorphs located in conserved but unlinked positions on pseudochromosomes 2 and 3, while only one MAT locus and idiomorph, *MAT1-1*, was identified in *Gh* on pseudochromosome 3 (Supplemental Fig. S8).

### Differences in effectors and secondary metabolite production potential between pathogenic and non-pathogenic *Gaeumannomyces* species

The predicted BGC content ranged from 9 to 17 per strain, which is low compared to many other ascomycete fungi (Gluck-Thaler et al. 2020; Franco et al. 2021; Llewellyn et al. 2023). Using a phylogenetically-informed permutational multivariate analysis of variance (PERMANOVA) method (Mesny and Vannier 2020) to identify associations between gene variance and lifestyle, we also found BGCs to have the lowest level of variance described purely by ancestry, 61% compared to 75%–85% for all genes, CSEPs and CAZymes (Fig. 4a). This was coupled with a relatively high proportion of BGC variance described by lifestyle (17%), which was also the case for all genes (17%) and CSEPs (14%), while lifestyle had much less descriptive power for CAZymes (7%). CAZymes that are known to act on plant cell wall substrates were highly conserved across the genus, and there were highly similar numbers of each CAZyme family across all strains (Supplemental Fig. S9a). The only discernible pattern was marginally more copies of GH55 and GH2 (hemicellulose and pectin) in *Gh* versus the other lineages.

In total, 9% of CSEP genes could be attributed to a known gene in the Pathogen-Host Interactions database (PHI-base) (Urban et al. 2020), most of which only had one copy in all strains (Supplemental Fig. S9b). Sixteen of the 19 ‘named’ CSEPs have been associated with virulence via reverse genetics experiments, including five from *P. oryzae* infecting *Oryza sativa* (rice) — *MHP1* (ID PHI:458); *MoAAT* (PHI:2144); *MoCDIP4* (PHI:3216); *MoHPX1* (PHI:5188); and *MoMAS3* (PHI:123065). The latter two were assigned to genes that were only present in *Gh*, although a separate gene present in *Gt*B was also characterised as *MoHPX1*. Six CSEPs in total were present in all lineages except *Gh* or vice versa. *PBC1*, also a CAZyme, the disruption of which causes complete loss of pathogenicity of *Pyrenopeziza brassicae* in *Brassica napus*, was present in *Gt* and *Ga* but not *Gh*. While *PBC1* was absent in *Gh*, all *Gaeumannomyces* strains did have some genes belonging to the same CAZyme family (CE5; Supplemental Fig. S9a).

The BGC families were predominantly classified as terpenes, type 1 polyketide synthases (PKSI) and hybrid polyketide synthase-nonribosomal peptides (PKS-NRP) (Supplemental Fig. S9c). Presence-absence of each BGC was highly variable across strains, most notably amongst PKSI families which also had high copy-number for certain families. There were five BGCs that were present or absent in *Gh* versus other lineages, four of which belonged to the PKS-NRP hybrids (Supplemental Fig. S9c).

### Gene copy-number reduction in *G. tritici* type A

*Gt*B, *Ga* and *Gh* all had high copy-number (HCN) gene outliers (>10 copies) that were absent in *Gt*A (Fig. 4a). These 22 HCN genes were duplicated both within and across pseudochromosomes (Supplemental Fig. S10a). GO term enrichment analyses found various terms to be significantly overrepresented amongst the HCN genes, namely: vacuolar proton-transporting V-type ATPase complex assembly (Gh-1B17, Fisher’s exact test, p=0.01); ubiquinone biosynthetic process (Gh-2C17, p=0.01); golgi organisation (Ga-CB1, p=0.03); mRNA cis splicing, via spliceosome (Gt-4e, p=0.03); mitochondrial respiratory chain complex I assembly (Gt-4e, p=0.05); proton-transporting ATP synthase complex assembly (Gt-LH10, p=0.03); and protein localisation to plasma membrane (Gt-LH10, p=0.03). Visualising the location of the HCN genes across the genomes (Supplemental Fig. S11) showed them to vary in terms of distribution — from relatively localised to broadly expanded — and in terms of multi-lineage versus lineage specific expansions. HCN genes were also significantly closer to TEs compared to other genes (Supplemental Fig. S10b).

Interestingly, of the 22 HCN genes, six that were shared among all species were also present in at least one *Gt*A strain but at low copy-number, while seven genes were completely absent in *Gt*A (Fig. 4c). In total, nine genes that were HCN in at least one other lineage had low-copy orthologues in *Gt*A. Moreover, these were mostly present in just one strain within the type A lineage (Gt-8d), clustered in a ∼1 Mbp region on pseudochromosome 3 (Supplemental Fig. S10c). This region was flanked by repetitive regions that have been subjected to repeat induced point mutation (RIP), as measured by the composite RIP index (CRI) (Lewis et al. 2009), although the region had average CRI of -0.3 compared to an average CRI of -0.5 for the whole pseudochromosome. Average genome-wide RIP levels were highest in *Gt*A and *Gh* (13.8% and 13.6% of the genome RIP’d, respectively), compared to *Gt*B (10.8%) and *Ga* (12.4%).

### *Gaeumannomyces* genomes contain *Starship* giant transposable elements

All nine *Gaeumannomyces* strains were found to contain at least one giant TE belonging to the *Starship* superfamily of giant cargo-carrying TEs (Gluck-Thaler et al. 2022), identified using the tool starfish (Gluck-Thaler and Vogan 2023). Currently the most reliable identifying feature of *Starships* is a single ‘captain’ gene – a tyrosine recombinase gene containing a DUF3435 domain which is found in the first position of each *Starship* and directs the mobilisation of the element (Urquhart et al. 2023b). We found that tyrosine recombinase annotation with starfish largely overlapped with results from a separate blast search to identify DUF3435 homologues at the head of insertions. Overall, only a relatively small number of genes were in agreement as full *Starship* captains after downstream automated (starfish) or manual element inference (Fig. 5a). A gene tree of all tyrosine recombinase and putative captain genes showed the presence of two distinct lineages but no consistent clustering of either gene types or method of identifying them. Note the highly divergent nature of the genes and therefore the difficulty of alignment and subsequent poor branch support throughout the tree (Fig. 5a).

**Figure 5.**
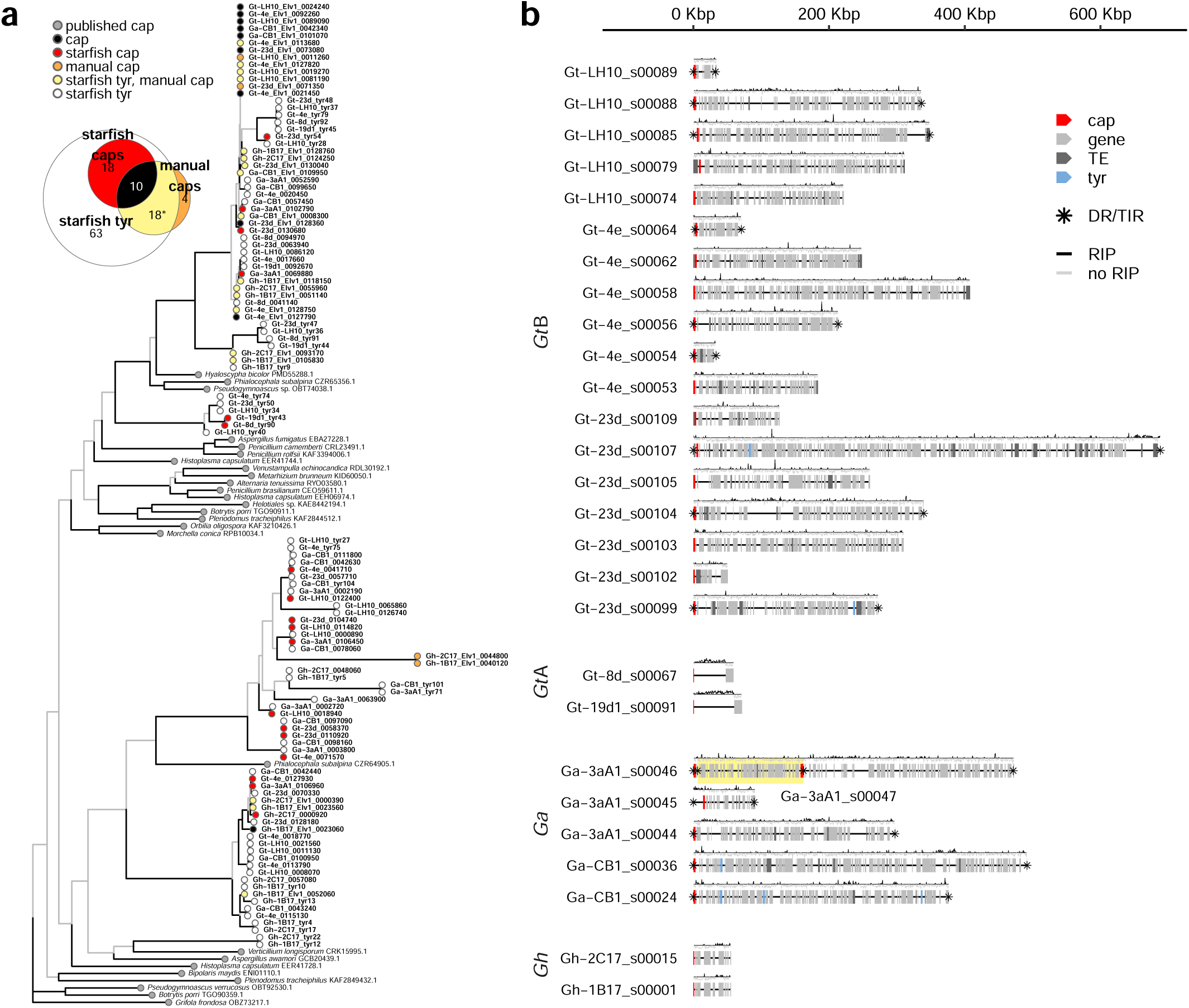
*Gaeumannomyces* genomes contain *Starship* giant transposable elements. **a** Gene tree of *Starship* ‘captain’ genes, including captains and other tyrosine recombinases identified from our assemblies via starfish, captain homologues identified via blastp, and previously published captain genes. **b** A summary of the *Starship* elements identified by starfish with the composite RIP index (CRI) shown above each element. The yellow highlight distinguishes a nested element. cap=captain gene, DR=direct repeat, RIP=repeat-induced point mutation, TE=transposable element gene, TIR=terminal inverted repeat, tyr=tyrosine recombinase gene.

*Starship* size varied considerably, ranging from 34–688 kbp. *Gt*B strains harboured notably more elements, followed by *Ga* strains which included a nested element (Fig. 5b). *Gt*A and *Gh* strains each contained a single smaller (<100 kbp) element, which in both cases we predict to have been vertically transmitted based on similar gene content and conserved location within the genome (Fig. 5b, Supplemental Fig. S12). *Gt*A elements were exceptional in that each was gene-poor and positive for element-wide RIP (average CRI=0.2-0.3).

## DISCUSSION

In this study we have established foundational genome resources for the genus *Gaeumannomyces*. A particular strength of the *Gt* assemblies reported here is the structural annotation methodology, which capitalised on the fact that multiple strains were sequenced, assembled and annotated in the same way, each with its own transcriptome but also employing a novel ‘multiple lift-off’ approach that provided additional evidence for robust gene models. Another benefit of the annotation approach is that the REAT-Mikado-minos pipeline (EI-CoreBioinformatics 2023b) provides models for gene isoforms alongside the primary transcripts. Alternative splicing has been implicated in regulation of virulence in phytopathogens (Fang et al. 2020), for instance by mediating transcriptome remodelling during pathogenesis in *P. oryzae* (Jeon et al. 2022). Alternative splicing has been reported to be more frequent in pathogens than non-pathogens (Grutzmann et al. 2014), however we found a similar overall percentage of genes with multiple isoforms in *Gh* compared to *Gt* and *Ga* (Supplemental Fig. S1). There was perhaps a skew towards a greater proportion of genes with exactly two or three isoforms in *Gt*, particularly *Gt*A, raising the question as to whether this somehow relates to their apparent higher virulence in wheat. These rich annotations resources will allow further exploration of the isoform content of *Gaeumannomyces* and its potential role in virulence.

A major finding from our synteny analyses was the presence of a large chromosomal translocation in Gt-LH10 (Fig. 2). Similar largescale translocations have been identified in *Pyricularia* (Bao et al. 2017; Gómez Luciano et al. 2019). It is entirely plausible that we have identified a genuine translocation, however confidence would be increased by obtaining Hi-C evidence and/or by corroborating with population-level data. Such rearrangements are thought to be a route to accessory chromosome formation (Croll et al. 2013; Hartmann 2022), and this has specifically been reported in *P. oryzae* (Langner et al. 2021). Although we did not find any evidence for accessory chromosomes in our *Gaeumannomyces* strains, it is interesting that the Gt-LH10 translocation resulted in one of the chromosomes being much smaller, size being a hallmark of accessory or ‘mini-chromosomes’. It is also notable that this large translocation occurred in the same strain we found to have an expansion of TEs (Supplemental Fig. S5), as TEs have been found to mediate interchromosomal rearrangements (Bao et al. 2017; Fourie et al. 2020; Langner et al. 2021). Hi-C data would also allow us to robustly locate centromeres (Varoquaux et al. 2015), which are also implicated in chromosomal rearrangements (Yadav et al. 2020; Guin et al. 2020). Here we used a minimal approach to estimate potential centromeric regions, based simply on the fact that AT-rich regions are a common defining feature of centromeres in *P. oryzae* (Yadav et al. 2019), which we also cross-checked with gene sparsity (Supplemental Fig. S2) — however, we were only able to distinguish potential centromeres for a subset of the pseudochromosomes.

In addition to the chromosomal translocation, Gt-LH10 also stood out from other strains in terms of TE content, with an expansion in total number of TEs (Supplemental Fig. S5) and smaller gene–TE distances (Fig. 3). Aside from the atypical features of the Gt-LH10 genome, there was additional intraspecific variability within the *Gt* A/B lineages in terms of both genome structure and gene content. For instance, there were strain-specific inversions (Fig. 2) and many of the HCN genes were present in low copy-number in one *Gt*A strain, but completely absent in the other (Fig. 4c). These findings emphasise the need for pangenome references, as an individual strain alone cannot sufficiently represent the variability across the whole species (Golicz et al. 2020; Badet and Croll 2020). Pangenomics is still relatively young and the practicalities of how to define, store and use pangenomes and the tools necessary to do so are continuously evolving (Eizenga et al. 2020). There is also the outstanding question of how best to coordinate pangenome initiatives to ensure high-quality results without duplicating efforts — at least three different pangenomes have been reported for the wheat pathogen *Zymoseptoria tritici* to date (Plissonneau et al. 2018; Badet et al. 2020; Chen et al. 2023). The five *Gt* strains reported here can act as a UK pangenome, but future research must work towards building a global pangenome so that we can provide a reference for *Gt* which captures a fuller representation of the species.

Another structural feature that these high-quality assemblies allowed us to explore in *Gaeumannomyces* was genome compartmentalisation. A number of fungal phytopathogens exhibit TE- and effector-rich compartments that enable rapid evolution in the plant–fungal arms race, dubbed the ‘two-speed’ genome model (Dong et al. 2015), which has since been extended to ‘multi-speed’ models (Frantzeskakis et al. 2019). Accordingly, we hypothesised that we would find CSEPs and TEs to colocalise across our assemblies, however we did not find consistent evidence for such compartments in *Gaeumannomyces* (Fig. 3). Our results are not altogether surprising as a previous study of selection signatures in *Gt* and two other *Magnaporthales* taxa also found no evidence for multi-speed genomes (Okagaki et al. 2016). We therefore consider *Gaeumannomyces* taxa to have ‘one-compartment’ genomes in relation to TE/effector content – a term that was introduced by Frantzeskakis et al. (2019) for genomes that do not conform to the two- or multi-speed models, and with ‘compartment’ suggested as an alternative to ‘speed’ as the defining features of these compartments does not necessarily equate to them being fast-evolving (Torres et al. 2020). With the rising number of high-quality genome resources, more examples are emerging that contradict the suggestion that phytopathogenicity is routinely accompanied by TE/effector compartmentalisation (Frantzeskakis et al. 2019). In fact, TE/effector compartmentalisation has been found in the non-pathogenic arbuscular mycorrhizal fungus *Rhizophagus irregularis* (Yildirir et al. 2022), and TE/virulence factor compartmentalisation has also been found in chytrid animal pathogens (Wacker et al. 2023), demonstrating that it is not necessarily central to phytopathogenicity, but may instead be a mechanism driving genome plasticity in fungi of various lifestyles (Torres et al. 2020). While we did not find compelling evidence for TE/effector compartmentalisation in *Gaeumannomyces*, we did observe non-random patterns in the distribution of CSEPs (Fig. 3a), which permutation analyses found to be closer to telomeric regions in a pseudochromosome-dependent manner (Supplemental Fig. S4b). This could suggest that alternative mechanisms of effector compartmentalisation may be at play.

Our results indicate conserved genetic machinery for plant cell wall deconstruction/ modification across both pathogenic and non-pathogenic *Gaeumannomyces* (Fig. 4a, S11a), suggesting that the mechanism(s) by which species first colonise roots may be similar, if not the final outcome of the plant-fungal interaction (Chancellor et al. 2023). Using spatial transcriptomics to visualise not only how *Gt* and *Gh* individually colonise wheat roots, but also how they interact with each other in the plant and the gene expression associated with this process, would undoubtedly shed light on this host-pathogen-antagonist system. Two putative orthologues of CSEP genes that have previously been implicated in pathogenicity were present in *Gt* and *Ga* pathogenic taxa but missing in non-pathogenic *Gh*, making them promising targets for future experiments to determine if either is important for *Gt* pathogenicity in wheat. *UvHrip1* (from *Ustilaginoidea virens*) is thought to be involved in suppressing host immunity and has already been reported in *Gt* (Wang et al. 2020), while *PBC1* (from *Pyrenopeziza brassicae*) is a cutinase implicated in host penetration (Li et al. 2003). It was intriguing that none of the CSEPs assigned to PHI-base genes were unique to *Gt*, perhaps suggesting that there is relatively high overlap in effector-mediated virulence mechanisms in *Gt* and *Ga*. In a similar pattern to the CSEPs, only one BGC (a PKS-NRP hybrid) was found to be specific to *Gt*, with most otherwise scattered across the genus (Supplemental Fig. S9c). There were more differences between *Gh* and the other lineages, and indeed the relative descriptive power of relatedness versus lifestyle on BGC variance (Fig. 4a) suggests that the production of secreted metabolites may be a key factor distinguishing the outcome of plant– fungal interactions in this genus. A single BGC has been shown to determine whether there is a mutualistic or pathogenic outcome of the interaction between root fungus *Colletotrichum tofieldiae* and *Arabidopsis thaliana* (Hiruma et al. 2023), demonstrating that minimal differences in metabolite repertoire can have considerable impacts on fungal lifestyle. In terms of host range, *Gt* has been shown to have low avenacinase activity relative to *Ga* (Osbourn et al. 1991), which is understood to be the reason *Gt* is incapable of also infecting oat roots (Osbourn et al. 1994). The avenacinase gene was nonetheless present in all strains across the genus; whether sequence polymorphism (Supplemental Fig. S7c) or differences in regulatory machinery are responsible for the variation in avenacinase activity remains to be determined. It is notable that *Gh* has also been found to be capable of colonising oat roots (Osborne et al. 2018) despite greater divergence of the *Gh* avenacinase protein sequence from *Ga* when compared to *Gt* (Supplemental Fig. S7b).

In line with the common understanding that *Gt* is self-fertile or homothallic (Palma-Guerrero et al. 2021), we found both *MAT1-1* and *MAT1-2* idiomorphs to be present in the *GtA* and *GtB* strains. These idiomorphs were located on two unlinked MAT loci, an atypical but occasionally observed homothallic MAT locus architecture in ascomycetes (Wilken et al. 2017; Dyer et al. 2016; Thynne et al. 2017). Although it is homothallic, *Gt* is also capable of outcrossing (Pilgeram and Henson 1992; Blanch et al. 1981), the rates of which may be underestimated in many other homothallic fungi (Billiard et al. 2012; Attanayake et al. 2014). Similarly to *Gt*, for *Ga* both MAT loci were identified. To our knowledge, the sex determination system of *Gh* has not previously been reported, but our results indicate only one idiomorph at a single MAT locus suggesting this species is self-sterile, or heterothallic. Evolutionary transitions between heterothallism and homothallism are common in ascomycetes (Thynne et al. 2017; Sun et al. 2019; Gioti et al. 2012; Ene and Bennett 2014), but the implications on fitness are not fully understood. In the scenario of a fungus infecting a crop monoculture, it may be advantageous for the fungus to be homothallic when rapidly expanding across the niche, as it will not be delayed by a reliance on the presence of compatible mating types. A higher rate of outcrossing due to heterothallism could be unfavourable, as it could break up combinations that are already well adapted to the genetically uniform host (Hill and McMullan 2023). Together, this could suggest a selective pressure towards homothallism for crop fungal pathogens, and a switch from heterothallism to homothallism may, therefore, have been a key innovation underlying lifestyle divergence between non-pathogenic *Gh* and pathogenic *Gt* and *Ga*.

An unanticipated result was the absence of HCN genes in the *Gt*A lineage (Fig. 4a), despite all other strains in the genus, including earlier diverging *Gh*, having genes which had undergone copy-number expansions (Supplemental Fig. S11). These HCN genes were on average significantly closer to TEs than other genes (Fig. 5c), which aligns with the fact that TEs are known to play a role in gene duplication (Cerbin and Jiang 2018). GO enrichment analysis identified a variety of fundamental biological processes to be significantly overrepresented amongst HCN genes in the other lineages: regulation of cellular pH and respiratory activity in non-pathogenic strains; and golgi organisation, protein localisation, mRNA cis-splicing and respiratory activity in pathogenic strains. As previously mentioned, alternative splicing has previously been linked to pathogenicity; respiratory activity has been shown to induce a developmental switch to symbiosis in an arbuscular mycorrhizal fungus (Tamasloukht et al. 2003); and mediation of cellular pH by V-ATPase has specifically been linked to pathogenesis in *P. oryzae* (Chen et al. 2013), although here it was implicated in a non-pathogenic *Gh* strain. Further investigation into the specific function of these genes is required to determine whether any of these processes are essential to lifestyle or virulence in *Gaeumannomyces*.

Gene duplicates are generally understood to be readily removed unless they serve to improve host fitness, for instance by favourably modifying expression levels or rendering a completely new function (Lynch and Conery 2000; Wapinski et al. 2007). RIP is a genome defence response against unchecked proliferation of duplicated sequences (Hane et al. 2015). In *Gaeumannomyces* we found 10–14% of the genome contained signatures of RIP, which is a moderate level relative to other ascomycetes, e.g. *Pyronema confluens* (0.5%) (Traeger et al. 2013), *Fusarium* spp. (<1–6%) (Van Wyk et al. 2019), *Neurospora* spp. (8–23%) (Gioti et al. 2013), *Zymoseptoria tritici* (14–35%) (Lorrain et al. 2021) and *Hymenoscyphus* spp. (24–41%) (Elfstrand et al. 2021). Genome-wide RIP was highest in *Gt*A, which was consistent with its low level of gene duplication, but not fully explanatory as *Gh* had only marginally lower levels of RIP while still maintaining HCN outliers. We can only presume that *Gt*A strains have been under stronger selective pressures to remove duplicates, although the evolutionary mechanisms driving this requires further investigation.

There was a similar pattern when exploring the RIP patterns across giant transposable *Starship* elements. We found only a single *Starship* in *Gt*A strains, which was gene-poor and had undergone extensive RIP (Fig. 5b), supporting the idea that this lineage employs stringent genome defence measures. By contrast, *Gt*B strains contained a proliferation of *Starships*, including one closely approaching the largest size reported thus far (Urquhart et al. 2023a). We expect that the increased availability of highly contiguous, long-read assemblies such as we report here will make the upper size extremes of such giant TEs more feasible to detect (Arkhipova and Yushenova 2019). Giant cargo-carrying TEs that can be both vertically and horizontally transmitted were first identified in bacteria (Johnson and Grossman 2015). Recently the *Starship* superfamily was identified as specific to and widespread in ascomycetes and, aside from the characteristic ‘captain’ tyrosine recombinase gene, each *Starship* contains a highly variable cargo (Gluck-Thaler et al. 2022). Mobilisation of cargo genes by *Starships* has been linked to the acquisition of various adaptive traits in fungal species, such as metal resistance (Urquhart et al. 2022), formaldehyde resistance (Urquhart et al. 2023a), virulence (McDonald et al. 2019), climatic adaptation (Tralamazza et al. 2023) and lifestyle switching (Gluck-Thaler et al. 2022). However, *Starships* are not inherently beneficial to the fungal host. One of the earliest groups of genes associated with the cargo of certain *Starships* was spore-killer or Spok genes, which bias their own transmission via the process of meiotic drive (i.e. by killing spores that do not inherit them) (Vogan et al. 2019). By incorporating Spok genes, a *Starship* element also biases its transmission, leading to it being referred to as a ‘genomic hyperparasite’ (Vogan et al. 2021). This corresponds to the concept of TEs as selfish genetic elements, which can prevail in the genome despite being neutral or deleterious to the overall fitness of the host. Whether mobilisation of an element and associated cargo is beneficial or detrimental to the host, TEs such as *Starships* are nonetheless drivers of genome evolution. Further detailed investigation of the specific cargo in the elements we have identified in *Gaeumannomyces* is a priority to explore how these giant TEs may be contributing to lifestyle and virulence.

While the differences in the overall appearance of the wheat plants and their root systems when infected with *Gt*A versus *Gt*B were visually compelling (Fig. 1A), our sample size was extremely limited and the quantitative data did not show such a strong distinction (Fig. 1C). A study by Lebreton et al. (2004) with a much larger sample size found *Gt* type A strains to be significantly more aggressive *in vitro* despite high intraspecific variability in take-all severity (type A=G2 in their study (Daval et al. 2010)). The dominance of type A strains in a site has also been reported to positively correlate with disease severity (Lebreton et al. 2007). It is also notable that five out of six wheat plants which died were inoculated with *Gt*A strains. Our phylogenomic analysis confirmed with significant branch support that the two lineages are indeed monophyletic (Supplemental Fig. S13b) and, together with our comparative genomics results, the question naturally arises as to whether *Gt*A and *Gt*B are in fact distinct species. We did not find evidence that genetic divergence between *Ga* and *Gt* species was more pronounced than between the *Gt*A and *Gt*B lineages, and host alone is not a sufficient distinction since, despite being a separate species, *Ga* is also able to infect wheat (Freeman and Ward 2004). Lebreton et al. (2004) suggested that ‘genetic exchanges between [A and B] groups are rare events or even do not exist’, but this was based on analysis of a limited number of genetic markers. Much broader whole-genome sequencing efforts are required to assess gene flow between lineages at the population-level, as well as the level of recombination. Understanding population dynamics could also shed light on the observed changes in ratio of *Gt*A and *Gt*B across wheat cropping years (Lebreton et al. 2004), which has implications for strategic crop protection measures.

### Conclusions

We have generated near-complete assemblies with robust annotations for under-explored but agriculturally important wheat-associated *Gaeumannomyces* species. In doing so we confirmed that *Gaeumannomyces* taxa have one-compartment genomes in the context of TE/effector colocalisation, however the presence of giant cargo-carrying *Starship* TEs likely contributes to genomic plasticity. Genomic signatures support the separation of *Gt* into two distinct lineages, with copy-number as a potential mechanism underlying differences in virulence. We also found evidence that variation amongst the relatively low number of BGCs may be a key factor contributing to lifestyle differences in *Gt* and *Gh*. In addition to providing foundational data to better understand this host–pathogen–antagonist system, these new resources are also an important step towards developing much-needed molecular diagnostics for take-all, whether conventional amplicon sequencing, rapid *in situ* assays (Hariharan and Prasannath 2021) or whole-genome/metagenomic sequencing approaches (Weisberg et al. 2021). Future research will require whole-genome sequencing of taxa from a broader geographical range to produce a global pangenome, which will provide a comprehensive reference for expression analyses to explore the role of virulence in *Gt* lineages, as well as population genomics to shed light on their evolution and distribution.

## METHODS

### Samples

Nine *Gaeumannomyces* strains were selected from the Rothamsted Research culture collections, including five *Gt* strains (two type A and three type B), two *Ga* strains and two *Gh* strains (Supplemental Table S2). All were collected from various experimental fields at Rothamsted Farm (Macdonald et al. 2018) between 2014 and 2018.

### *G. tritici* virulence test in adult wheat plants

To test the virulence of the five *Gt* strains, we performed inoculations of each strain (six replicates) into the highly susceptible winter wheat cultivar Hereward. First the roots of seedling plants were inoculated with the fungus by using plastic drinking cups (7.5 cm wide x 11 cm tall) as pots, ensuring that all seedlings were well colonised before transferring to a larger pot. Pots were drilled with four drainage holes 3 mm in diameter. A 50 cm^3^ layer of damp sand was added to each pot, followed by a 275 g layer of naïve soil collected from a field at Rothamsted Farm after a non-legume break crop. Inoculum was prepared by taking a 9 mm fungal plug with a cork borer number 6 from the outer part of a fungal colony grown on a potato dextrose agar (PDA) plate and mixing with sand to make up a 25 g inoculum layer. A final 150 g layer of naïve soil was added on top of the inoculum layer. One wheat seed was sown on the surface of the soil and covered with a 50 cm^3^ layer of grit to aid germination and create a humid environment for fungal colonisation. Pots were watered well and placed in a controlled environment room (16 hr day, light intensity 250 μmols, 15°C day, 10°C night, watered twice a week from above). A randomised block design was generated in Genstat 20th Edition to take potential environmental differences across the growth room into account.

After two weeks of growth, each wheat seedling in a small pot was transferred by removing the plastic cup and placing the entire contents undisturbed into a larger 20 cm diameter pot containing a 2 cm layer of clay drainage pebbles. Three small pots were transferred to each large pot and filled in with more soil, resulting in three plants per pot. There were 6 replicates for each treatment, and a control pot with no fungus was also set up in the same manner, but a PDA plate without fungus was used for preparing the inoculum layer. The pots were transferred to a screenhouse and arranged randomly within blocks containing one pot per treatment. The pots were established in September and remained outside in the screenhouse to ensure exposure to winter conditions and therefore allow plant vernalisation to take place.

Measurements of the above-ground characteristics were first undertaken to note the severity of any take-all symptoms once the floral spike (ear) was fully emerged. The height of each labelled plant was measured from the stem base to the tip of the ear to the nearest 0.5 cm to identify whether there was stunted growth. Additionally, the length of the ear and flag leaf were recorded, again to the nearest 0.5 cm. The number of ears per plant was also recorded.

For below-ground measurements, the pots were washed out post full plant senescence and the plants were well rinsed to remove the soil while minimising damage to the roots. Any roots that broke off were collected and put into the cup with the main plants to maintain accuracy of the biomass measurements. The stems were then cut about 10 cm from the base. The plants were placed in a white tray filled with water to enable clear observation of the roots. The number of tillers for each plant was counted. The severity of take-all infection was then estimated by using the Take-All Index (TAI), classified through the following categories: Slight 1 (0–10% of roots infected), slight 2 (11–25%), moderate 1 (25–50%), moderate 2 (51–75%) and severe (76–100%). This was then input into the following formula: TAI = ((1 x % plants slight 1) + (2 x % plants slight 2) + (3 x % plants moderate 1) + (4 x % plants moderate 2) + (5 x % plants severe)) / 5 (Bateman et al. 2004). Following this, the length of the roots was measured to the nearest 0.5 cm. By cutting off one root at a time, the number of roots for each plant was counted and the roots transferred into cardboard trays, one per pot. These were then dried at 80°C on metal trays for 16 hours. One tray at a time was removed from the oven to reduce any moisture gain before weighing. The dried root biomass per pot was then recorded.

To statistically test for mean differences in the various characteristics between strains, we first made Q-Q plots using the ggqqplot function from ggpubr v0.6.0 (Kassambara 2020) to confirm approximate data normality. We then used the levene_test function from the package rstatix v0.7.2 (Kassambara 2021) to assess the assumption of homogeneity of variance, where a significant p value (p < 0.05) means that the assumption is violated. If we could ascertain homogeneity of variance, a multiple comparison test between strains was performed with the tukey_hsd rstatix function. Where homogeneity of variance was violated, the games_howell_test rstatix function was instead used for multiple comparison testing (Sauder and DeMars 2019).

### Genome sequencing

For DNA and RNA extractions of all nine *Gaeumannomyces* taxa, a 4 mm plug of mycelium from axenic cultures was transferred to 500 ml of potato dextrose broth treated with penicillium/streptomycin (10,000 U/mL) using a sterile 4 mm corer. Cultures were grown at 20°C in dark conditions on an orbital shaker at 140 rpm for ∼ 7–14 days. Mycelia were collected via vacuum filtration and flash frozen using liquid nitrogen and stored at -80°C, before grinding with a sterilised mortar and pestle until a fine powder was created.

DNA was extracted using one of two kits: the Phytopure Nucleon Genomic DNA kit (Cytiva, MA, USA) eluted in 50 µl low-pH TE buffer; and the NucleoBond HMW DNA kit (Macherey-Nagel, North Rhine-Westphalia, Germany) eluted in 100 µl–200 µl low-pH TE buffer. The manufacturer’s protocols were modified to optimise for high molecular weight (M. Grey, personal communication). Sufficient DNA concentration (50 ng/µl DNA) was confirmed by Qubit fluorometer (Invitrogen, MA, USA) and purity (260/280 absorbance ratio of approximately 1.6–2.0 and 260/230 absorbance ratio of approximately 1.8–2.4) confirmed with a NanoDrop spectrophotometer (Thermo Fisher Scientific, MA, USA). Sufficient strand lengths (80% > 40 Kbp length) were confirmed using the Femto Pulse System (Agilent Technologies, Inc, CA, USA).

RNA from the same sample material was extracted using the Quick-RNA Fungal/Bacterial miniprep kit (Zymo Research, CA, USA) using the manufacturer’s protocol and eluted in 25 µl of DNase/RNase free water. Sufficient RNA concentration (71 ng/µl RNA) was confirmed by Qubit fluorometer (Invitrogen, MA, USA) and purity (260/280 absorbance ratio of approximately 1.8–2.1 and 260/230 absorbance ratio of > 2.0) confirmed with a NanoDrop spectrophotometer (Thermo Fisher Scientific, MA, USA). An RNA integrity number > 8 was confirmed by Bioanalyzer RNA analysis (Agilent Technologies, Inc, CA, USA).

DNA and RNA extractions were sent to the Genomics Pipelines Group (Earlham Institute, Norwich, UK) for library preparation and sequencing. 2–5.5 µg of each sample was sheared using the Megaruptor 3 instrument (Diagenode, Liege, Belgium) at 18-20ng/µl and speed setting 31. Each sample underwent AMPure PB bead (PacBio, CA, USA) purification and concentration before undergoing library preparation using the SMRTbell Express Template Prep Kit 2.0 (PacBio) and barcoded using barcoded overhang adapters 8A/B (PacBio) and nuclease treated with SMRTbell enzyme cleanup kit 1.0 (PacBio). The resulting libraries were quantified by fluorescence (Invitrogen Qubit 3.0) and library size was estimated from a smear analysis performed on the Femto Pulse System (Agilent). The libraries were equimolar pooled into four multiplex pools and each pool was size fractionated using the SageELF system (Sage Science, MA, USA), 0.75% cassette (Sage Science). The resulting fractions were quantified by fluorescence via Qubit and size estimated from a smear analysis performed on the Femto Pulse System, and 1–2 fractions per pool were selected for sequencing and pooled equimolar to have equal representation of barcodes per pool. The loading calculations for sequencing were completed using the PacBio SMRTLink Binding Calculator v10.1.0.119528 or v10.2.0.133424. Sequencing primer v2 or v5 was annealed to the adapter sequence of the library pools. Binding of the library pools to the sequencing polymerase was completed using Sequel II Binding Kit v2.0 or 2.2 (PacBio). Calculations for primer to template and polymerase to template binding ratios were kept at default values. Sequel II DNA internal control was spiked into the library pool complexes at the standard concentration prior to sequencing. The sequencing chemistry used was Sequel II Sequencing Plate 2.0 (PacBio) and the Instrument Control Software v10.1.0.119549 or 10.1.0.125432. Each pool was sequenced on 1-2 Sequel II SMRTcells 8M (PacBio) on the Sequel IIe instrument. The parameters for sequencing were as follows: CCS sequencing mode; 30-hour movie; 2-hour adaptive loading set to 0.85 or diffusion loading; 2-hour immobilisation time; 2–4-hour pre-extension time; and 70–86pM on plate loading concentration.

RNA libraries were constructed using the NEBNext Ultra II RNA Library prep for Illumina kit (New England Biolabs, MA, USA), NEBNext Poly(A) mRNA Magnetic Isolation Module and NEBNext Multiplex Oligos for Illumina (96 Unique Dual Index Primer Pairs) at a concentration of 10 µM. RNA libraries were equimolar pooled, q-PCR was performed, and the pool was sequenced on the Illumina NovaSeq 6000 (Illumina, CA, USA) on one lane of a NVS300S4 flowcell with v1.5 chemistry producing a total of 3,370,873,981 reads.

### Genome assembly

See Supplemental Fig. S14a for a schematic summarising the bioinformatics analyses. HiFi reads were assembled using hifiasm v0.16.1-r375 (Cheng et al. 2021) with the -l 0 option to disable purging of duplicates in these haploid assemblies. The assemblies were checked for content correctness with respect to the input HiFi reads using the COMP tool from KAT v2.3.4 (Mapleson et al. 2017), and QUAST v5.0.2 (Mikheenko et al. 2018) was used to calculate contiguity statistics. BlobTools v1.0.1 (Laetsch and Blaxter 2017) was used to check for contamination (Supplemental Fig. S15) — this required a hits file, which we produced by searching contigs against the nt database (downloaded 21/05/2021) using blastn v2.10, and a BAM file of mapped HiFi reads, which we produced using minimap2 v2.21 (Li 2018) and samtools v1.13 (Li et al. 2009).

Gene set completeness was assessed using the ascomycota_odb10.2020-09-10 dataset in BUSCO v5.2.1 (Manni et al. 2021). This revealed some gene duplication due to the presence of small contigs that had exceptionally low coverage (median of 1 across each small sequence) when projecting the kmer spectra of the reads onto them using KAT’s SECT tool. This was taken as evidence that the sequences did not belong in the assemblies. A custom script was written to filter out these small, low-coverage sequences, using the output of KAT SECT. KAT COMP, BUSCO and QUAST were re-run for the coverage filtered assemblies to verify that duplicated genes were removed without losing core gene content and produce final assembly contiguity statistics (Supplemental Fig. S16, Supplemental Table S1).

### Genome annotation

Repeats were identified and masked using RepeatModeler v1.0.11 (Smit and Hubley 2015) and RepeatMasker v4.0.7 (Smit et al. 2015) via EIRepeat v1.1.0 (Kaithakottil and Swarbreck 2023). Gene models were annotated via the Robust and Extendable Eukaryotic Annotation Toolkit (REAT) v0.6.3 (EI-CoreBioinformatics 2023b) and MINOS v1.9 (EI-CoreBioinformatics 2023a). The REAT workflow consists of three submodules: transcriptome, homology, and prediction. The transcriptome module utilised Illumina RNA-Seq data, reads that were mapped to the genome with HISAT2 v2.1.0 (Kim et al. 2019) and high-confidence splice junctions identified by Portcullis v1.2.4 (Mapleson et al. 2018). The aligned reads were assembled for each tissue with StringTie2 v1.3.3 (Kovaka et al. 2019) and Scallop v0.10.2 (Shao and Kingsford 2017). A filtered set of non-redundant gene models were derived from the combined set of RNA-Seq assemblies using Mikado v2.3.4 (Venturini et al. 2018). The REAT homology workflow was used to generate gene models based on alignment of protein sequences from publicly available annotations of 27 related species (Supplemental Table S3) and a set of proteins downloaded from UniProt including all the proteins from the class *Sordariomycetes* (taxid:147550) and excluding all proteins from the publicly available annotation of *Gt* R3-111a-1 (GCF_000145635). The prediction module generated evidence-guided models based on transcriptome and proteins alignments using AUGUSTUS v3.4.0 (Stanke et al. 2006), with four alternative configurations and weightings of evidence, and EVidenceModeler v1.1.1 (Haas et al. 2008). In addition, gene models from the *Gt* R3-111a-1 annotation were projected via Liftoff v1.5.1 (Shumate and Salzberg 2021), and filtered via the multicompare script from the ei-liftover pipeline (Venturini and Yanes 2020), ensuring only models with consistent gene structures between the original and transferred models were retained.

The filtered Liftoff, REAT transcriptome, homology and prediction gene models were used in MINOS to generate a consolidated gene set with models selected based on evidence support and their intrinsic features. Confidence and biotype classification was determined for all gene models based on available evidence, such as homology support and expression. TE gene classification was based on overlap with identified repeats (> 40 bp repeat overlap).

To make best use of having multiple identically generated annotations for the genus, we opted to additionally repeat a lift-over process projecting the gene models from each MINOS run to all nine assemblies. We then removed gene models overlapping rRNA genes from the multiple-lift-over annotations and the previously consolidated MINOS annotation using RNAmmer v1.2 (Lagesen et al. 2007) and BEDTools v2.28 (Quinlan and Hall 2010). The MINOS consolidation stage was repeated using four files as input: the high-confidence models from the lift-over; the high-confidence genes of the previous MINOS run for the specific assembly; the low-confidence models of the previous MINOS run for the specific assembly; and the low-confidence models of the lift-over of all the closely related species. This multiple-lift-over approach allowed us to cross-check gene sets across strains and determine whether missing genes were truly absent from individual assemblies or had just been missed by the annotation process. Finally, mitochondrial contigs were identified using the MitoHiFi v2.14.2 pipeline (Uliano-Silva et al. 2023), with gene annotation using MitoFinder v1.4.1 (Allio et al. 2020) and the mitochondrion sequence from *Epichloë novae-zelandiae* AL0725 as a reference (GenBank accession NC_072722.1).

Functional annotation of the gene models was performed using AHRD v3.3.3 (Hallab et al. 2023), with evidence from blastp v2.6.0 searches against the Swiss-Prot and TrEMBL databases (both downloaded on 19/10/2022), and mapping of domain names using InterProScan v5.22.61 (Jones et al. 2014). Additional annotations were produced using eggNOG-mapper v2.1.9 (Cantalapiedra et al. 2021) with sequence searches against the eggNOG orthology database (Huerta-Cepas et al. 2019) using DIAMOND v2.0.9 (Buchfink et al. 2021). CAZymes were predicted using run_dbcan v3.0.1 (Le and Yohe 2021) from the dbCAN2 CAZyme annotation server (Zhang et al. 2018) this process involved (i) HMMER v3.3.2 (Mistry et al. 2013) search against the dbCAN HMM (hidden Markov model) database; (ii) DIAMOND v2.0.14 search against the CAZy pre-annotated CAZyme sequence database (Drula et al. 2022) and (iii) eCAMI (Xu et al. 2020) search against a CAZyme short peptide library for classification and motif identification. A gene was classified as a CAZyme if all three methods were in agreement.

CSEPs were predicted using a similar approach to Hill et al. (2022), with some additions/substitutions of tools informed by Jones et al. (2021); see Supplemental Fig. S14b for a schematic overview. Briefly, we integrated evidence from SignalP v3.0 (Dyrløv Bendtsen et al. 2004), v4.1g (Petersen et al. 2011), v6.0g (Teufel et al. 2022); TargetP v2.0 (Almagro Armenteros et al. 2019); DeepSig v1.2.5 (Savojardo et al. 2018); Phobius v1.01 (Käll et al. 2004); TMHMM v2.0c (Krogh et al. 2001); Deeploc v1.0 (Almagro Armenteros et al. 2017); ps_scan v1.86 (Gattiker et al. 2002); and EffectorP v1.0 (Sperschneider et al. 2016), v2.0 (Sperschneider et al. 2018) and v3.0 (Sperschneider and Dodds 2021). CSEPs were then matched to experimentally verified genes in the PHI-base database (Urban et al. 2020) (downloaded 21/07/2023) using a BLAST v2.10 blastp search with an e-value cutoff of 1e-25. In the event of multiple successful hits, the hit with the top bitscore was used. Secondary metabolites were predicted using antiSMASH v6.1.1 (Blin et al. 2021). Reference protein sequences for avenacinase from *Ga* (GenBank accession AAB09777.1) and mating-type locus idiomorphs *MAT1-1* and *MAT1-2* from *Pyricularia grisea* (Latorre et al. 2022) were used to identify their respective genes in each of the nine assemblies using a blastp search (e-value cutoff 1e-25).

### Phylogenetic classification of *G. tritici* types

To confirm the classification of *Gt* strains within established genetic groups — *sensu* Daval et al. (2010) and Freeman et al. (Freeman et al. 2005) — gene trees were produced for gentisate 1,2-dioxygenase (*gdo*; GenBank accessions FJ717712–FJ717728) and *ITS2*. GenePull (Hill 2021) was used to extract the two marker sequences from the new assemblies reported here. *ITS2* could not be found in the existing *Gt* R3-111a-1 assembly (RefSeq accession GCF_000145635.1), so that strain was only included in the *gdo* gene tree. We aligned each marker gene separately using MAFFT v7.271 (Katoh and Standley 2013) and manually checked the gene alignments. The gene trees were estimated using RAxML-NG v1.1.0 (Kozlov et al. 2019) and the GTR+G nucleotide substitution model (Supplemental Fig. S13a). Branch support was computed using 1,000 Felsenstein’s bootstrap replicates, or until convergence according to the default 3% cutoff for weighted Robinson-Foulds distances (Pattengale et al. 2009), whichever occurred first. An avenacinase gene tree was produced in the same way but using the JTT+G4 amino acid substitution model.

### Phylogenomics of *Gaeumannomyces*

A genome-scale species tree was produced to provide evolutionary context for comparative analyses. We used OrthoFinder v2.5.4 (Emms and Kelly 2019) to cluster predicted gene models for primary transcripts into orthogroups — in addition to the newly sequenced *Gaeumannomyces* taxa, this also included *Gt* R3-111a-1 and the outgroup *Magnaporthiopsis poae* ATCC 64411 (GenBank accession GCA_000193285.1). Alongside the coalescent species tree produced within OrthoFinder by STAG (Emms and Kelly 2018), we also used a concatenation-based approach. We used MAFFT to produce gene alignments for 7,029 single-copy phylogenetic hierarchical orthogroups or HOGs (hereafter, genes) that were present in all taxa. These were trimmed using v1.4.rev15 (Capella-Gutiérrez et al. 2009), concatenated using AMAS and run in RAxML-NG with genes partitioned and the JTT+G4 amino acid substitution model. Branch support was calculated as above.

Alongside the species tree we visualised assembly N50; the number of gene models; the proportion of these that were functionally annotated by AHRD; and the number of unassigned gene models from OrthoFinder (Supplemental Fig. S17). Due to concerns regarding the comparability of the existing *Gt* R3-111a-1 annotation to the strains reported in this study, and to avoid introducing computational bias, the existing *Gt* R3-111a-1 annotation was excluded from downstream comparative analyses for the sake of consistency.

### Genome structure and synteny

To identify both potential misassemblies and real structural novelty in our strains, we used GENESPACE v1.1.8 (Lovell et al. 2022) to visualise syntenic blocks across the genomes. Fragments were considered to have telomeres at the ends if Tapestry v1.0.0 (Davey et al. 2020) identified at least five telomeric repeats (TTAGGG), and this was used together with the GENESPACE results to inform pseudochromosome designation. Telomeric repeats were also cross-checked with results from tidk v0.2.31 (Brown 2023). We calculated GC content across pseudochromosomes in 100,000 bp windows using BEDTools v2.29.2 (Quinlan and Hall 2010), and TE, gene and CSEP density were calculated in 100,000 bp windows with a custom script, plot_ideograms.R. The composite RIP index (CRI) (Lewis et al. 2009) was calculated in 500 bp windows using RIP_index_calculation.pl (Stajich 2023).

To statistically test for correlations between CSEP density and TE and /or gene density, we again made Q-Q plots using the ggqqplot function to assess approximate data normality. This being violated, we calculated Kendall’s tau for each strain (rstatix cor_test function, method="kendall"). The assumption of normality being similarly violated for distances from CSEPs/other genes to the closest TE, we performed a Wilcoxon rank sum test (wilcox_test function) to compare mean distances for CSEPs versus other genes for each strain. To compare the mean gene–TE distance across strains, we used a Games-Howell test (games_howell_test function) for multiple comparison testing. Comparison of distances between HCN genes and TEs versus other genes and TEs was tested in the same way.

We also performed permutation tests of CSEP–TE distances using the meanDistance evaluation function from the R package regioneR v1.32.0 (Gel et al. 2016), with the resampleRegions function used for randomisation of the gene universe over 1,000 permutations. Permutation tests of CSEP–telomere distances were performed in the same way, having assigned the first and last 10,000 bp of each pseudochromosome as telomeric regions.

### Comparative genomics

Functional annotations were mapped to orthogroups using a custom script, orthogroup_assigner.R, adapted from Hill et al. (2022), which also involved retrieval of CAZyme names from the ExplorEnz website (McDonald et al. 2009) using the package rvest v1.0.3 (Wickham 2020). CAZyme families known to act on the major plant cell wall substrates were classified as by Hill et al. (2022) based on the literature (Glass et al. 2013; Levasseur et al. 2013; Zhao et al. 2013; Miyauchi et al. 2020; Hage and Rosso 2021; Mesny et al. 2021). For *Gt*, gene content was categorised as core (present in all strains), soft-core (present in all but one strain), accessory (present in at least two strains) and specific (present in one strain).

Broadscale differences in gene repertoires due to lifestyle (pathogenic *Gt* and *Ga* and non-pathogenic *Gh*) were statistically tested using a permutational analysis of variance (PERMANOVA) approach to estimate residual variance of gene content after accounting for variance explained by phylogenetic distance (Mesny and Vannier 2020). To analyse the potential for secondary metabolite production with this PERMANOVA approach, a presence-absence matrix for biosynthetic gene cluster families was produced from the antiSMASH results using BiG-SCAPE v1.1.5 (Navarro-Muñoz et al. 2020).

Gene duplicates were categorised as intrachromosomal (on the same pseudochromosome) or interchromosomal (on a different pseudochromosome) using the pangenes output files from GENESPACE. We conducted gene ontology (GO) enrichment analysis for high copy-number (HCN) genes using the R package topGO v2.50.0 (Alexa and Rahnenfuhrer 2022) with Fisher’s exact test and the weight01 algorithm.

### *Starship* element identification

Giant transposable *Starship* elements were identified in our assemblies after noting dense blocks of transposons forming gaps between annotated genes. Manual inspection of these regions via synteny plots built with OMA v2.5.0 (Altenhoff et al. 2019) and Circos v0.69 (Krzywinski et al. 2009) revealed *Starship*-sized insertions (Gluck-Thaler et al. 2022), and an NCBI blastp search of the first gene in one such insertion in strain Gt-8d (Gt-8d_EIv1_0041140) returned 85% identity with an established *Gt* R3-111a-1 DUF3435 gene (GenBank accession EJT80010.1). These two genes were then used for a local blastp v2.13.0 search against all nine *Gaeumannomyces* assemblies reported here, which identified 33 full length hits (>95% identity) that were associated with insertions when visualised in Circos plots. This manual approach was then compared to *Starship* element identification using starfish v1.0 (Gluck-Thaler and Vogan 2023). One element identified by starfish was discounted as it consisted solely of a single predicted captain gene with no cargo or flanking repeats. A gene tree of all tyrosine recombinases predicted by starfish (including *Starship* captains), blastp-identified DUF3435 homologues, and previously reported *Starship* captain genes (Gluck-Thaler et al. 2022) was built using the same methods described above for phylogenetic classification and the JTT+G4 amino acid substitution model, with the addition of alignment trimming using trimAl v1.4.rev15 (Capella-Gutiérrez et al. 2009) with the -gappyout parameter. Data visualisation was completed in R v4.3.1 (R Core Team 2022) using the packages ape v5.7-1 (Paradis and Schliep 2019), aplot v0.2.2 (Yu et al. 2023), ComplexUpset v1.3.3 (Krassowski 2022), cowplot v1.1.1 (Wilke 2020), data.table v1.14.8 (Dowle and Srinivasan 2023), eulerr v7.0.0 (Larsson 2020), ggforce v0.4.1 (Pedersen 2021), ggh4x v0.2.6 (van den Brand 2023), gggenomes v0.9.12.9000 (Hackl et al. 2023), ggmsa v1.6.0 (Zhou et al. 2022), ggnewscale v0.4.9 (Campitelli 2020), ggplot2 v3.4.4 (Wickham 2016), ggplotify v0.1.2 (Yu 2021), ggpubr v0.6.0 (Kassambara 2020), ggrepel v0.9.3 (Slowikowski 2020), ggtree v3.9.1 (Yu et al. 2017), Gviz v1.44.2 (Hahne and Ivannek 2016), matrixStats v1.0.0 (Bengtsson 2021), multcompView v0.1-9 (Graves et al. 2019), patchwork v1.1.3 (Pederson 2022), rtracklayer v1.60.1 (Lawrence et al. 2009), scales v1.2.1 (Wickham and Seidel 2020), seqmagick v0.1.6 (Yu 2023), tidyverse v2.0.0 (Wickham et al. 2019). All analysis scripts are available at https://github.com/Rowena-h/GaeumannomycesGenomics.

## DATA ACCESS

WGS data and annotated genome assemblies are available on GenBank under the BioProject accession PRJNA935249 (assemblies pending release). Additional data files are deposited in Zenodo doi:10.5281/zenodo.10277622 (pending release). All bioinformatics scripts are available at https://github.com/Rowena-h/GaeumannomycesGenomics.

## ACKNOWLEDGEMENTS

We thank Jonathan Storkey at Rothamsted Research for grass species identification following the sampling of the Park Grass long term pasture experiment at Rothamsted. Many thanks to Suzanne Clark in the Computational and Analytical Sciences group at Rothamsted Research for the randomised block design in the inoculation experiments and for the subsequent data analyses. Thanks to Rachel Rusholme Pilcher for help and advice with GENESPACE, and other members of the Anthony Hall lab group for general discussion. The authors acknowledge funding from the Biotechnology and Biological Sciences Research Council (BBSRC), part of UK Research and Innovation, Core Capability Grants BB/CCG1720/1 and the work delivered via the Scientific Computing group, as well as support for the physical HPC infrastructure and data centre delivered via the NBI Computing infrastructure for Science (CiS) group. This research was funded by the BBSRC grants Designing Future Wheat (BB/P016855/1) and Delivering Sustainable Wheat (BB/X011003/1) and the constituent work packages (BBS/E/ER/230003B Earlham Institute and BBS/E/RH/230001B Rothamsted Research). Part of this work was delivered via the BBSRC National Capability in Genomics and Single Cell Analysis (BBS/E/T/000PR9816) at the Earlham Institute by Suzanne Henderson of the Genomics Pipelines Group. The long-term Park Grass field experiment at Rothamsted Research and the entire Rothamsted Experimental Farm is supported by the UK Biotechnology and Biological Sciences Research Council (BBSRC) and the Lawes Agricultural Trust. D Smith is supported by the BBSRC Core Capability Grant (BB/CCG2280/1). GC was supported by the DEFRA funded Wheat Genetic Improvement Network (WGIN) phase 3 (CH0106) and GC and JS were supported by WGIN phase 4 (CH0109). S-JO and TC were supported by the BBSRC funded University of Nottingham Doctoral Training Programme (BB/M008770/1). JH was supported by a UK government– Rothamsted Research level 3 Laboratory Technician Apprenticeship scheme.

## Author contributions

RH, MM, NH, KH-K and JP-G conceived, managed and/or coordinated the work. NH, MM and KH-K were involved in funding acquisition. MM, JPG and KH-K supervised different aspects of the research. GC, VEM, S-JO, JH, MG and MM collected the samples and/or isolated strains. JP-G, JS and JH performed *Gt* inoculation experiments. MG and NI performed molecular lab work. SJW performed genome assembly analyses. MOF performed genome annotation analyses, supervised by D Swarbreck. RH performed functional annotation analyses; designed and performed phylogenetic, comparative and statistical analyses; and performed data visualisations. D Smith performed the exploratory *Starship* analyses. RH and MM wrote the manuscript with contributions from MG, MOF, TC, D Smith, NI, NH, JP-G, and KH-K. All authors read and approved the manuscript.

