## Supplementary Material for "Evolutionary genomics reveals variation in structure and genetic content implicated in virulence and lifestyle in the genus *Gaeumannomyces*"

*Authors for correspondence:

**Table S1** Statistics for the new assemblies reported in this study.

| | QUAST v5.0.2 | | | | | BUSCO v5.2.1 (n=1,706) | MitoHifi v2.14.2 | | |
|---|---|---|---|---|---|---|---|---|---|
| | # contigs ≥ 500bp | Largest contig (bp) | Total size (bp) | GC (%) | N50 | Assembly completeness (single-copy BUSCOs) | Mitogenome size (bp) | GC (%) | # genes |
| *Gaeumannomyces avenae* Ga-CB1 | 37 | 7,079,356 | 41,409,597 | 56.93 | 4,890,083 | 1,654 (97.0%) | 167,275 | 31.41 | 39 |
| *G. avenae* Ga-3aA1 | 62 | 9,242,662 | 41,681,452 | 56.77 | 6,143,224 | 1,654 (97.0%) | 172,445 | 31.57 | 39 |
| *G. hyphopodioides* Gh-1B17 | 69 | 8,478,668 | 48,861,344 | 57.25 | 6,286,456 | 1,587 (93.0%) | 113,602 | 30.70 | 40 |
| *G. hyphopodioides* Gh-2C17 | 49 | 8,472,645 | 45,138,018 | 57.46 | 6,996,108 | 1,656 (97.1%) | 116,357 | 30.72 | 40 |
| *G. tritici* Gt-8d | 41 | 9,558,045 | 41,248,712 | 56.54 | 6,032,443 | 1,652 (96.8%) | 174,693 | 31.59 | 38 |
| *G. tritici* Gt-19d1 | 63 | 8,750,995 | 41,352,696 | 56.35 | 6,009,945 | 1,664 (97.5%) | 174,693 | 31.59 | 38 |
| *G. tritici* Gt-23d | 54 | 9,793,078 | 44,534,174 | 57.05 | 6,017,529 | 1,656 (97.1%) | 190,005 | 31.42 | 38 |
| *G. tritici* Gt-4e | 45 | 10,701,577 | 43,603,089 | 57.31 | 6,461,149 | 1,662 (97.4%) | 182,019 | 31.42 | 38 |
| *G. tritici* Gt-LH10 | 332 | 15,505,531 | 52,080,895 | 54.08 | 5,872,440 | 1,662 (97.4%) | 60,839 | 31.17 | 23 |

**Table S2** Metadata for the *Gaeumannomyces* strains sequenced in this study. *Soil cores collected post-grain harvest of a wheat plot.

| **Species** | **Strain** | **Host/substrate** | **Collection year** | **Rothamsted Research field** |
|---|---|---|---|---|
| *Gaeumannomyces avenae* | Ga-CB1 | Common bent (*Agrostis capillaris*) | 2018 | Park Grass |
| *G. avenae* | Ga-3aA1 | Soil | 2018 | Park Grass |
| *G. hyphopodioides* | Gh-1B17 | Wheat (*Triticum aestivum*, Hereward) | 2016 | New Zealand |
| *G. hyphopodioides* | Gh-2C17 | Wheat (*Triticum aestivum*, Scout) | 2016 | New Zealand |
| *G. tritici* | Gt-8d | Soil* | 2017 | Long Hoos |
| *G. tritici* | Gt-19d1 | Soil* | 2017 | Long Hoos |
| *G. tritici* | Gt-23d | Soil* | 2017 | Long Hoos |
| *G. tritici* | Gt-4e | Soil* | 2017 | Long Hoos |
| *G. tritici* | Gt-LH10 | Soil* | 2014 | Long Hoos |

**Table S3** NCBI accessions for assemblies which were used to inform structural annotation.

| Strain | Accession |
|---|---|
| *Magnaporthiopsis poae* ATCC 64411 | GCA_000193285 |
| *Valsa mali* 03-8 | GCA_000818155 |
| *Epichloë festucae* Fl1 | GCA_003814445 |
| *Pyricularia* sp. CBS 133598 | GCA_004337975 |
| *Fusarium* culmorum Class2-1B | GCA_016952355 |
| *Trichoderma simmonsii* GH-Sj1 | GCA_019565615 |
| *Trichoderma semiorbis* FJ059 | GCA_020045945 |
| *Trichoderma cornu-damae* KA19-0412C | GCA_020631695 |
| *Fusarium solani-melongenae* CRI 24-3 | GCA_023101225 |
| *Colletotrichum lupini* IMI 504893 | GCA_023278565 |
| *Pyricularia oryzae* 70-15 | GCF_000002495 |
| *Neurospora crassa* OR74A | GCF_000182925 |
| *Fusarium verticillioides* 7600 | GCF_000149555 |
| *Thermothelomyces thermophilus* ATCC 42464 | GCF_000226095 |
| *Thermothielavioides terrestris* NRRL 8126 | GCF_000226115 |
| *Fusarium graminearum* PH-1 | GCF_000240135 |
| *Fusarium pseudograminearum* CS3096 FP7 | GCF_000303195 |
| *Ustilaginoidea virens* UV-8b | GCF_000687475 |
| *Drechmeria coniospora* ARSEF 6962 | GCF_001625195 |
| *Pochonia chlamydosporia* 170 | GCF_001653235 |
| *Colletotrichum higginsianum* IMI 349063 | GCF_001672515 |
| *Pyricularia pennisetigena* PpBr36 | GCF_004337985 |
| *Pyricularia grisea* PgNI | GCF_004355905 |
| *Fusarium poae* DAOMC 252244 | GCF_019609905 |
| *Fusarium musae* F31 | GCF_019915245 |
| *Purpureocillium takamizusanense* PT3 | GCF_022605165 |
| *Fusarium venenatum* A3/5 | GCF_900007375 |

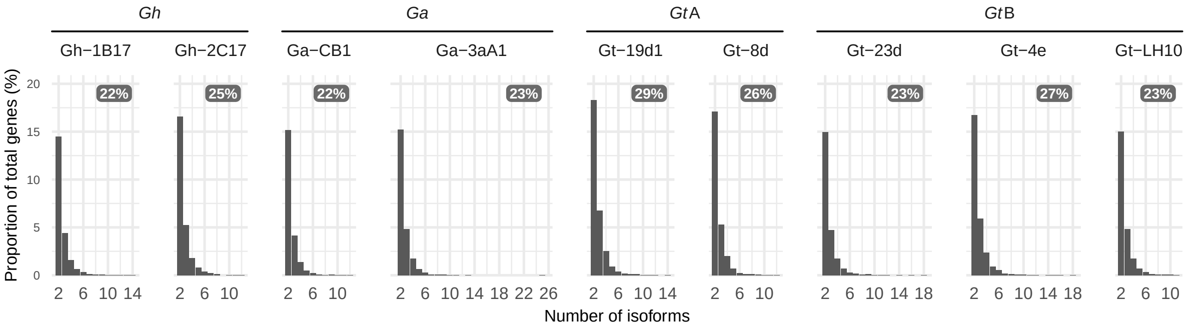


**Fig. S1** Histograms showing the proportion of genes with multiple isoforms annotated by REAT. Overall percentage of genes with two or more isoforms are shown in boxes to the top right of each plot.

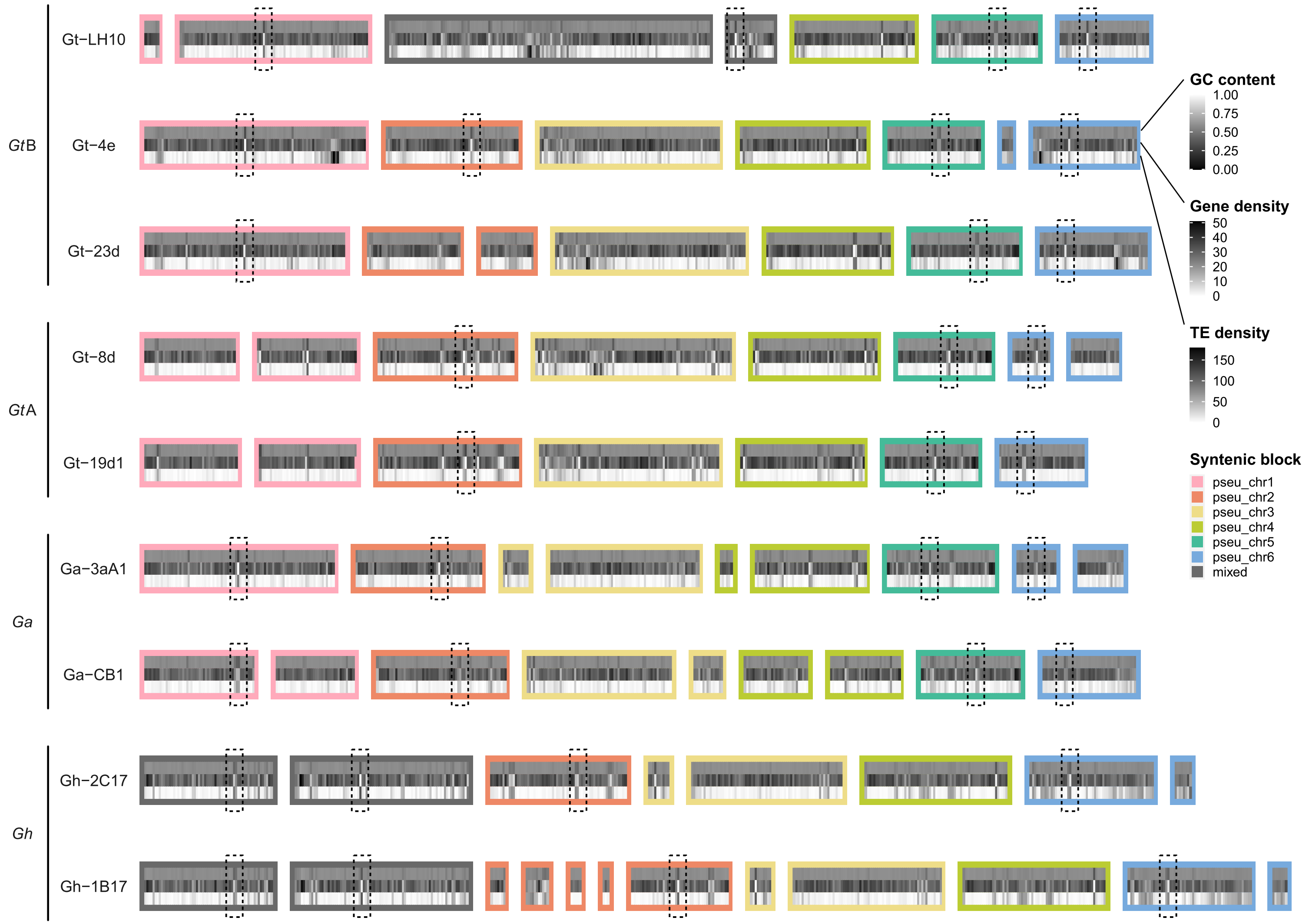


**Fig. S2** Putative centromeres identified from GC content, transposable element (TE) density and gene density. All calculations were done in 100,000 bp windows.

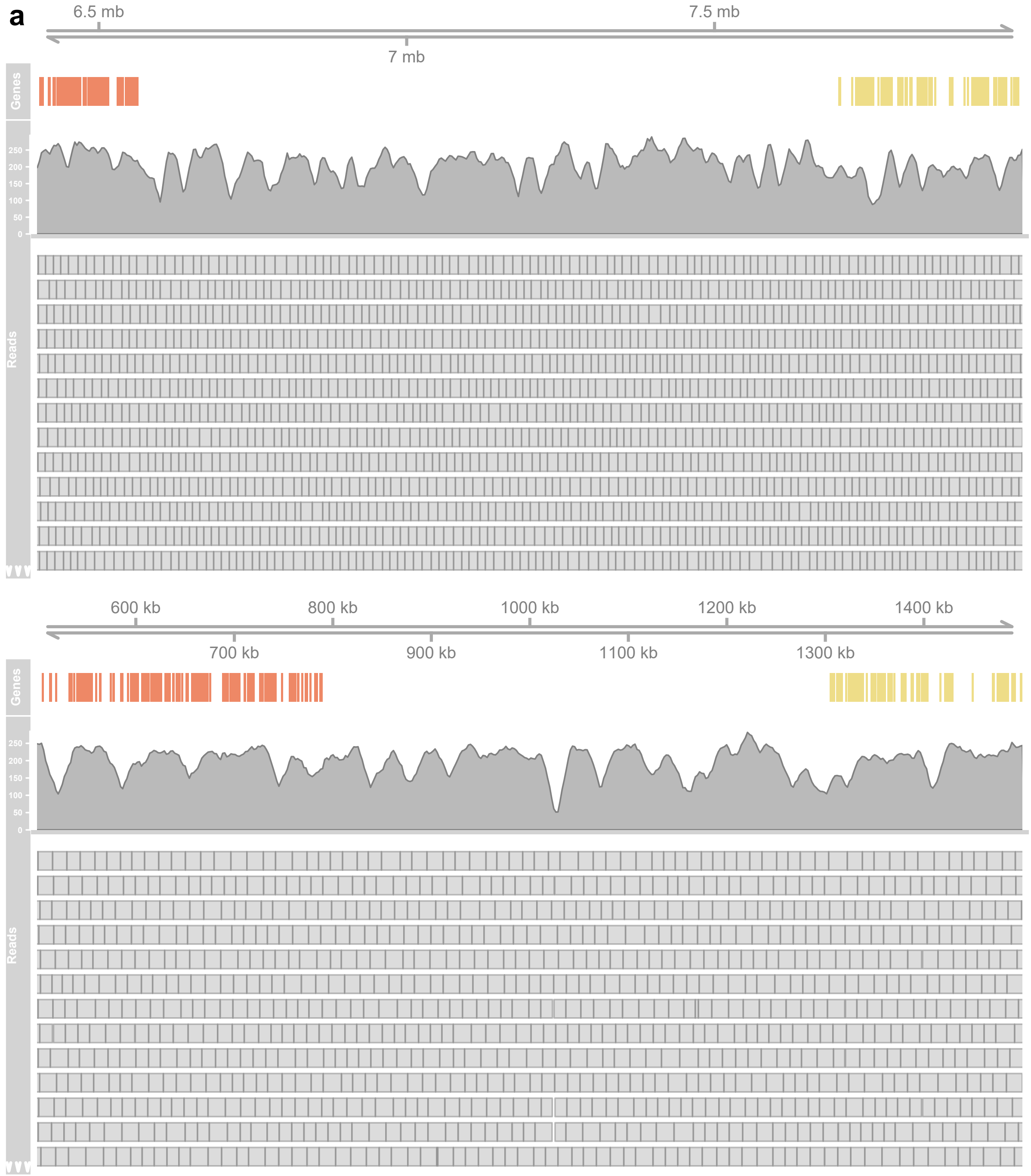


**Fig. S3 a** Read coverage across the putative translocation in pseudochromosome 2B (top) and pseudochromosome 3B (bottom) in strain Gt-LH10. Grey boxes indicate the location of reads mapped to the assembly. Gene colours match the GENESPACE syntenic blocks in Fig. 2. Note that pseudochromosome 3B has not been inverted as in the GENESPACE plot. ▼

b

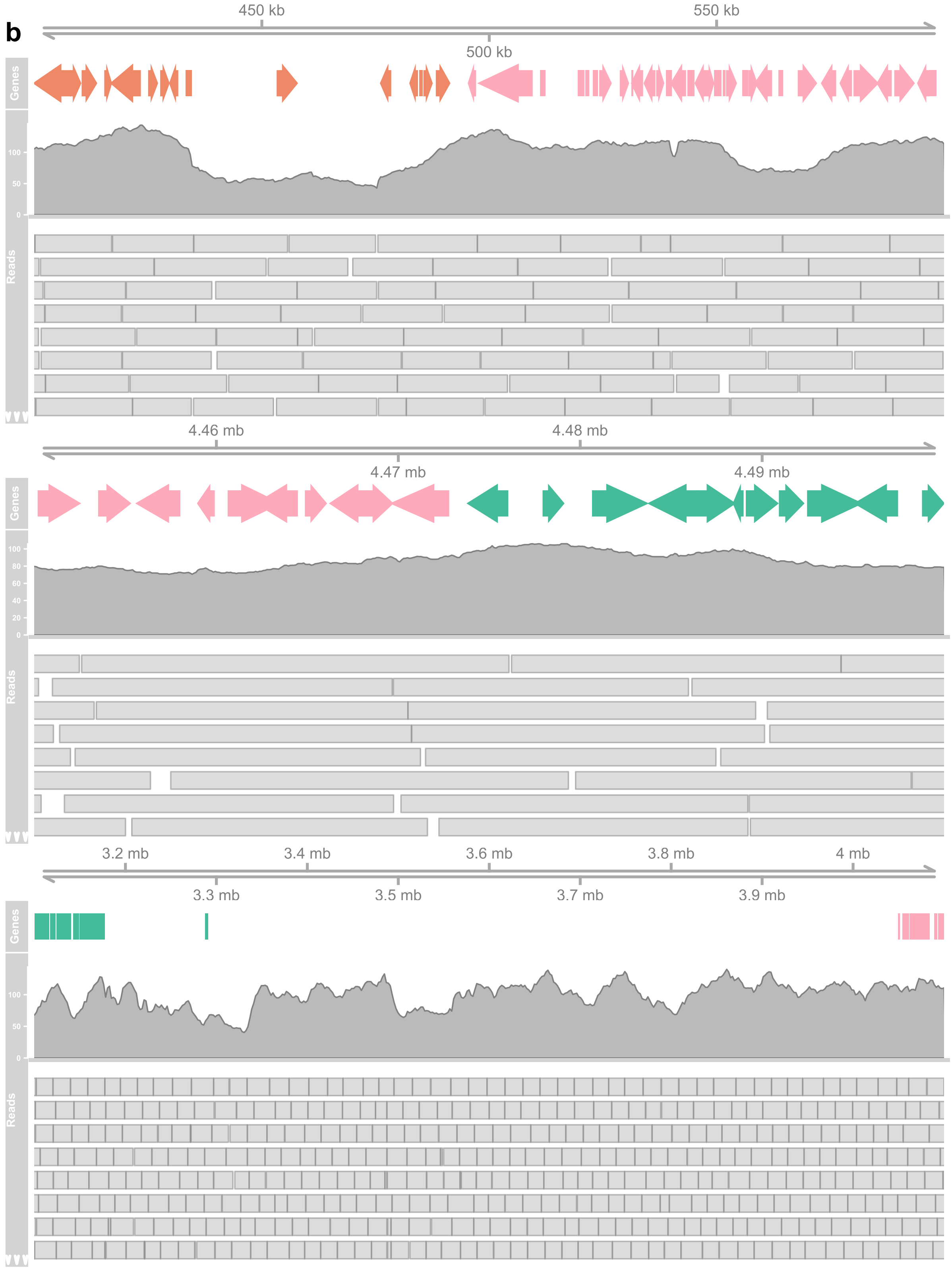


**Fig. S3 continued b** Read coverage across the putative translocations in pseudochromosome 1B (top and middle) and pseudochromosome 5B (bottom) in strain Gh-1B17. Grey boxes indicate the location of reads mapped to the assembly. Gene colours match the GENESPACE syntenic blocks in Fig. 2. ▼

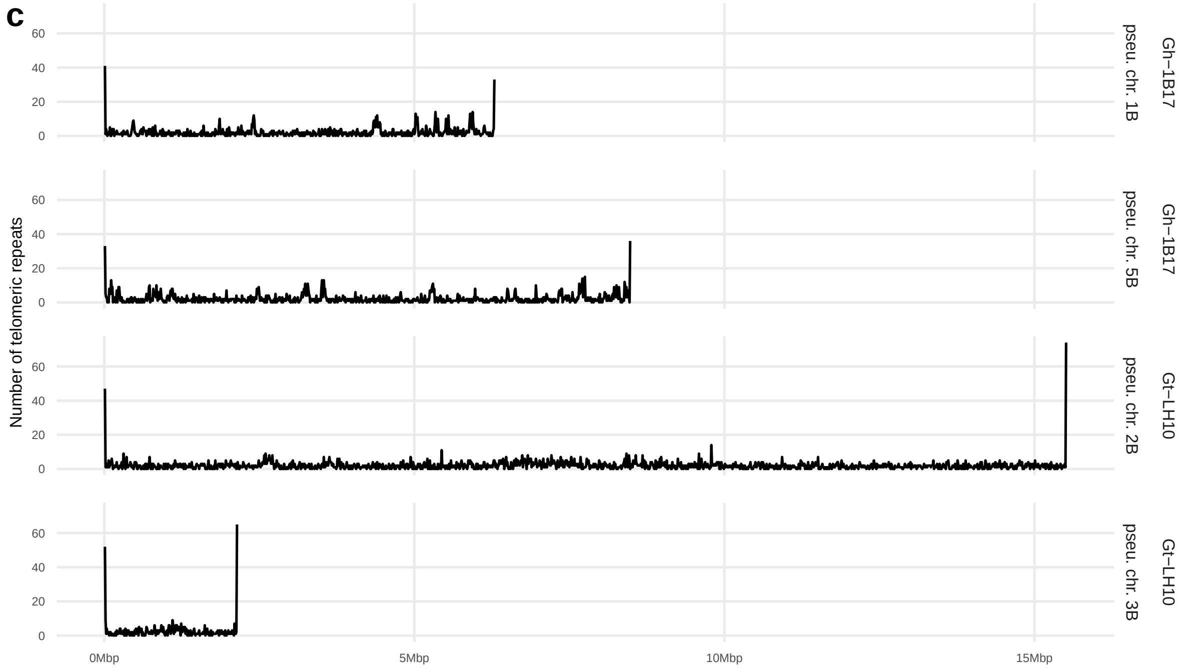


**Fig. S3 continued c** Telomeric repeats as estimated by tidk across Gh-1B17 pseudochromosomes 1B and 5B and Gt-LH10 pseudochromosomes 2B and 3B.

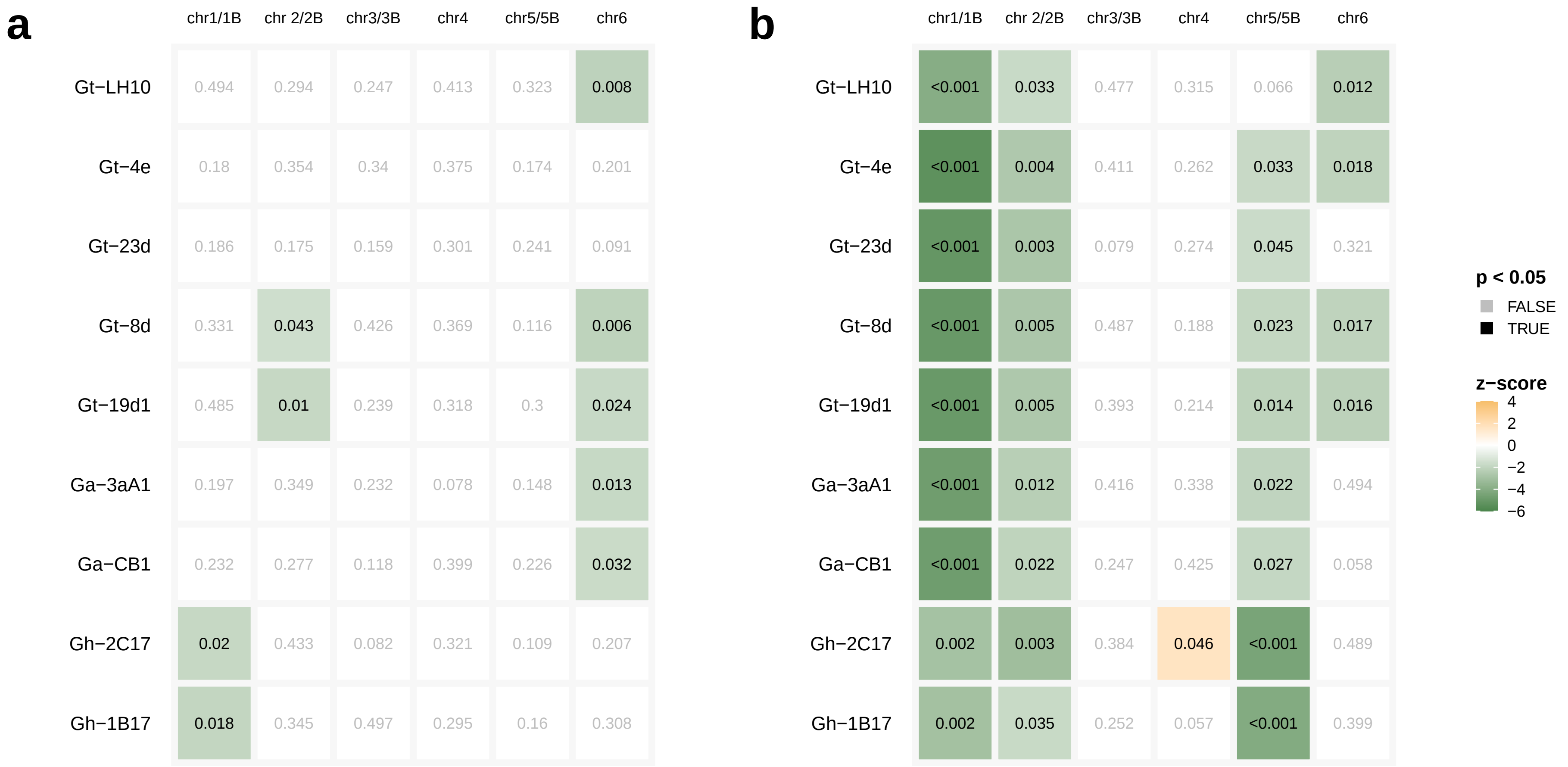


**Fig. S4** Matrices of p values from permutation analyses showing whether candidate secreted effector proteins (CSEPs) are significantly closer than expected to **a** transposable elements (TEs) and **b** telomeres. Each test used 1,000 permutations with random resampling of the gene universe for the pseudochromosome. Statistically significant test results ($p < 0.05$) are coloured by z-score, a proxy for 'strength' of the test result, where negative z-score means smaller than expected distance and positive means greater than expected.

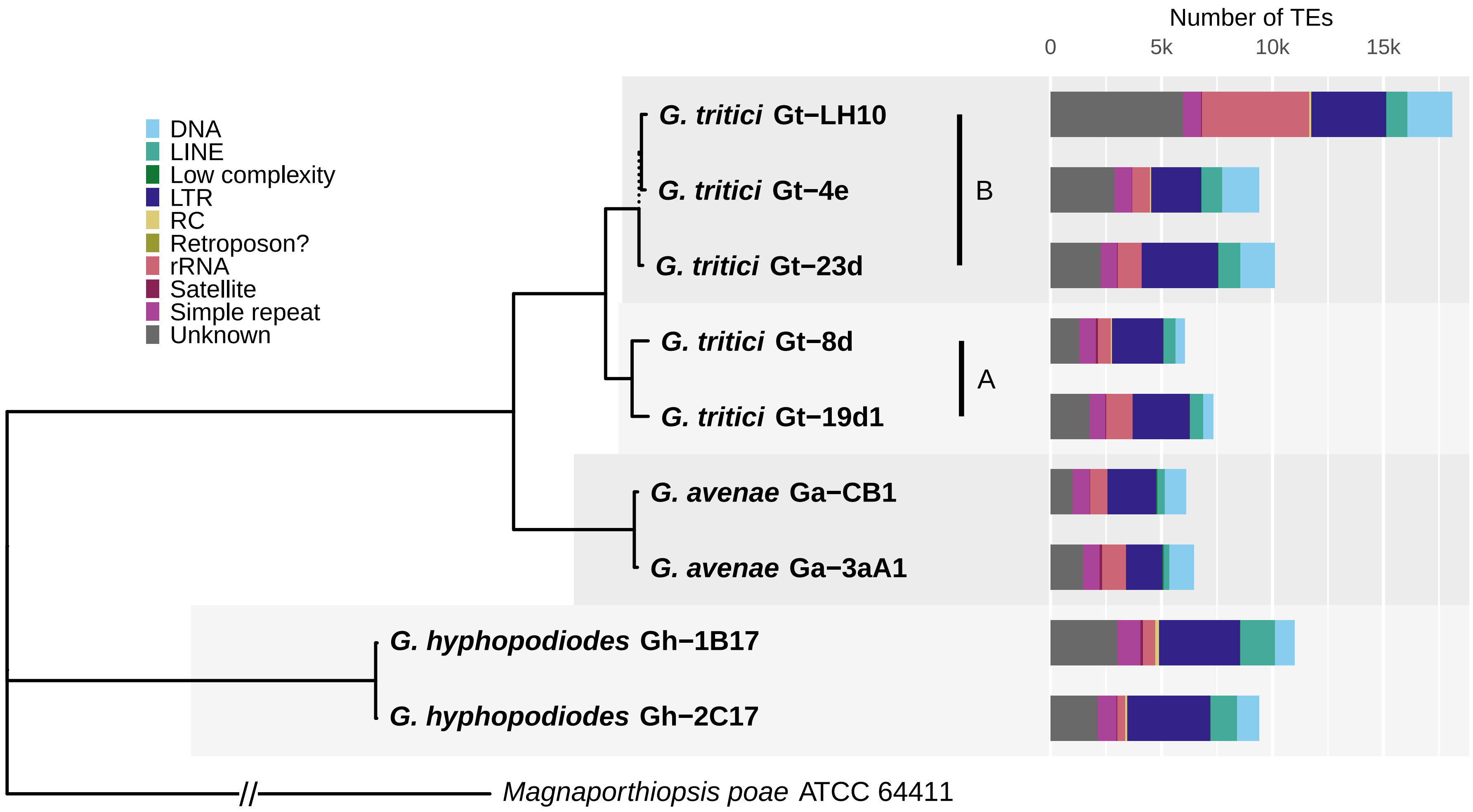


**Fig. S5** Total number of transposable elements (TEs) for each strain and their classification as assigned by RepeatMasker.

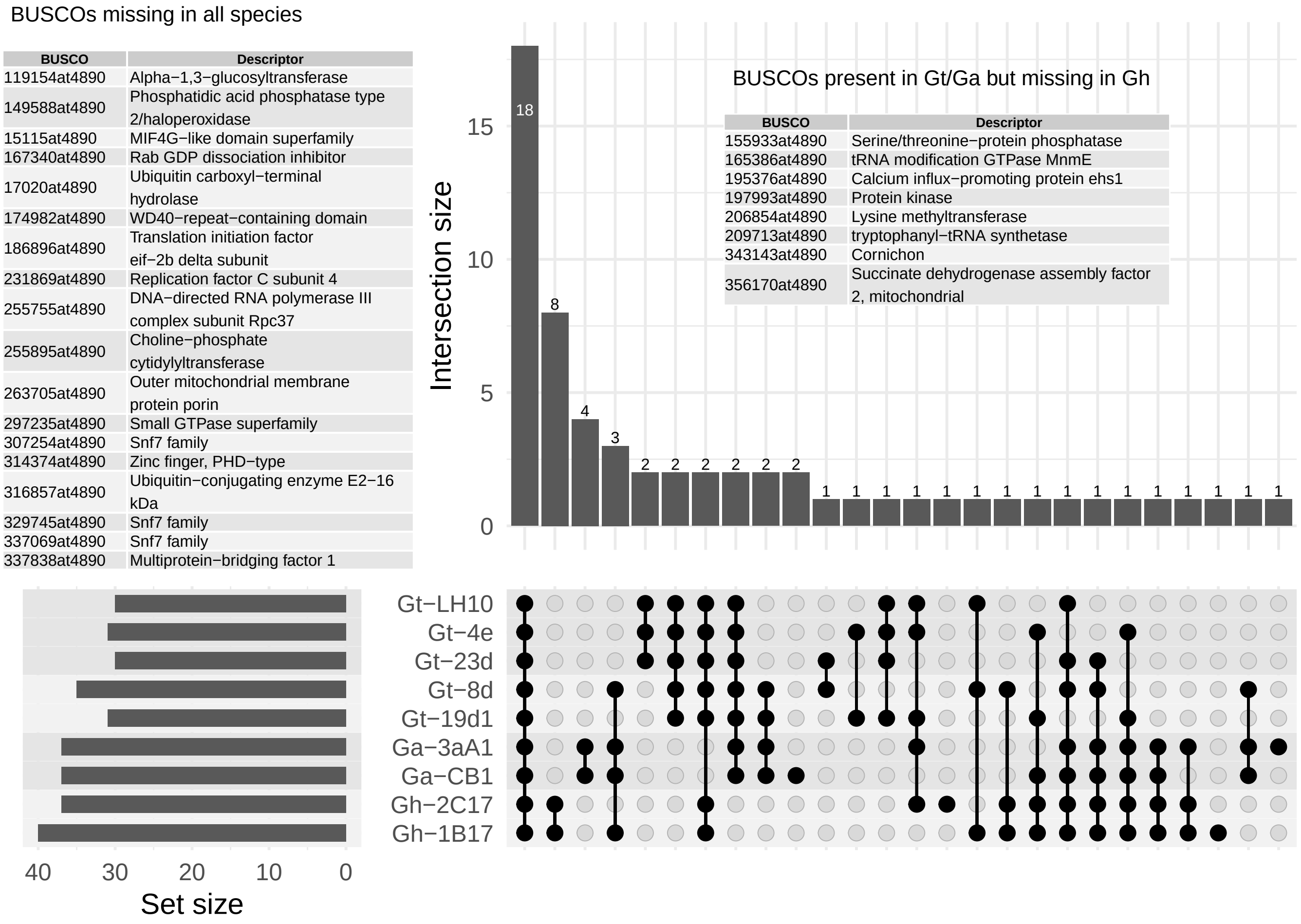


**Fig. S6** Upset plot showing the intersections of missing BUSCOs across the *Gaeumannomyces* assemblies. Inset tables list specific BUSCOs missing in all species and missing in just *G. hyphopodioides*.

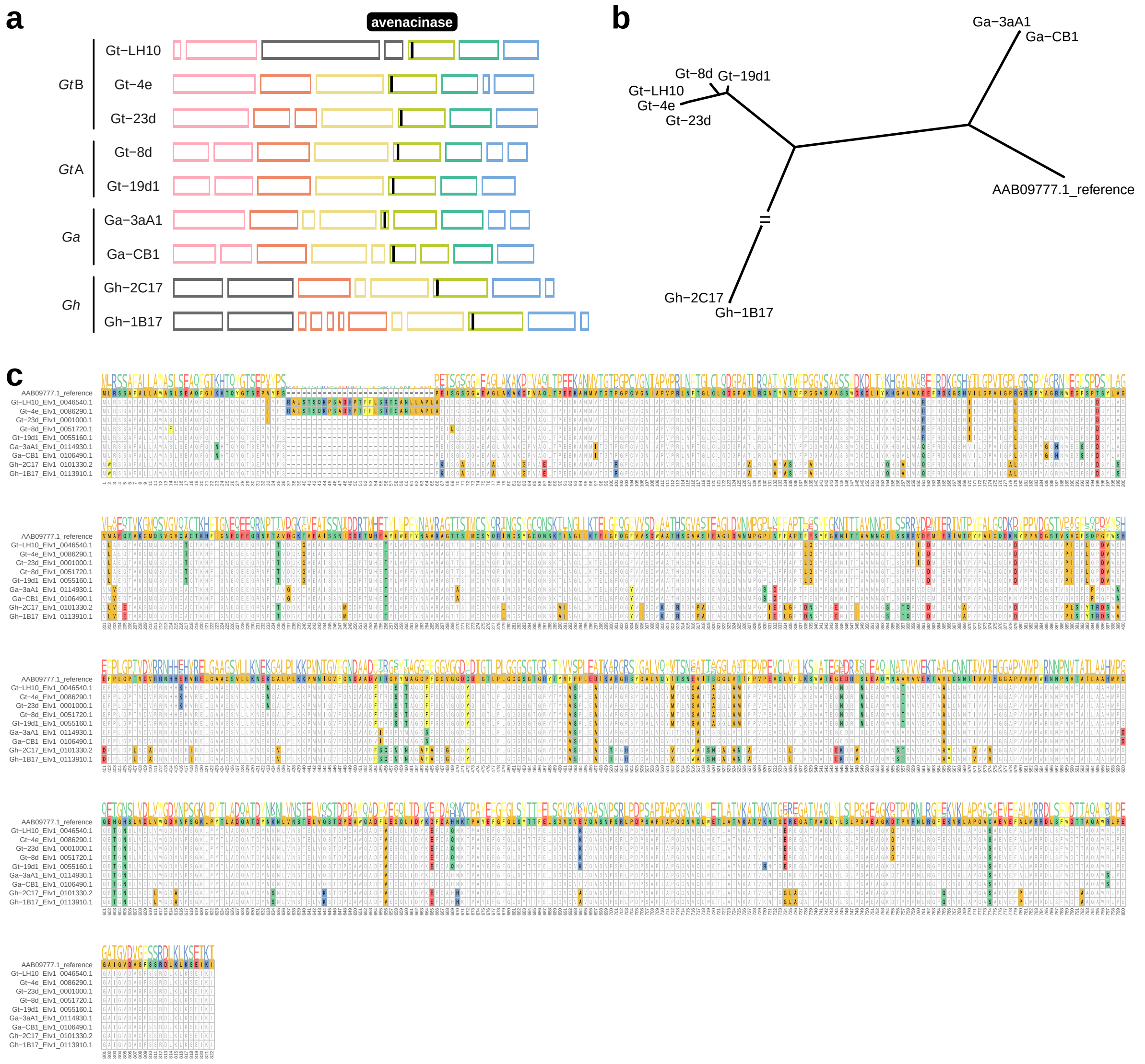


**Fig. S7 a** Conserved location of the avenacinase gene on pseudochromosome 4. **b** An unrooted RAxML-NG gene tree of the avenacinase gene. **c** Protein sequence alignment of the avenacinase gene, with amino acids coloured if they disagree with the reference sequence in the top row.

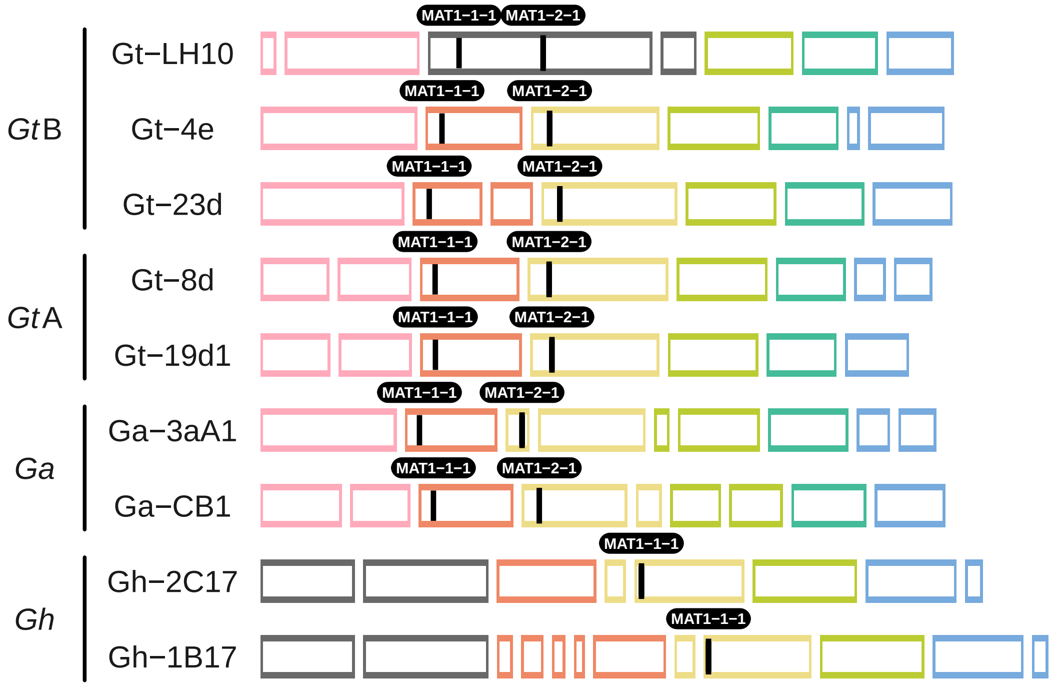


**Fig. S8** Location of MAT loci.

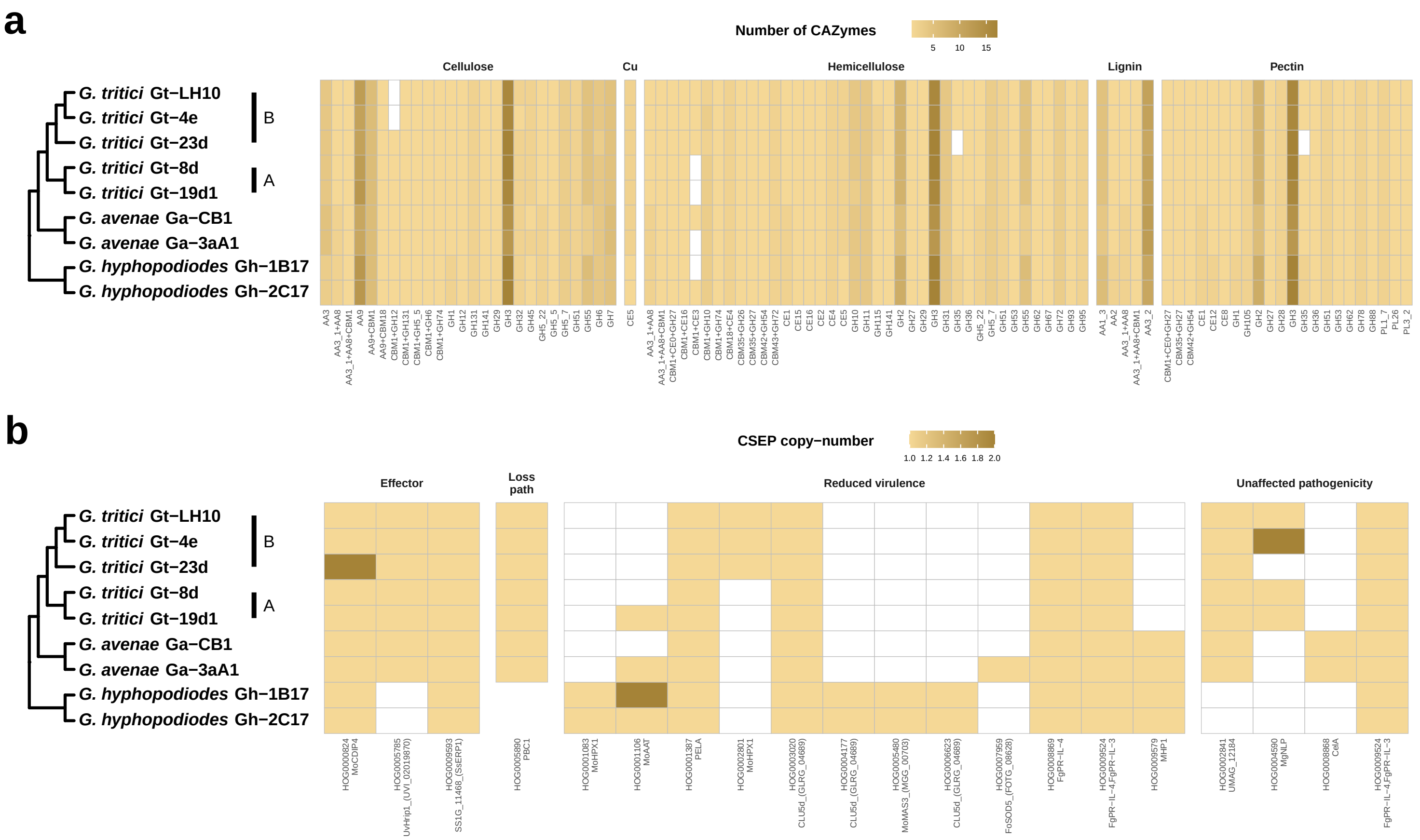

**Fig. S9** **a** Abundance matrix showing number of genes belonging to carbohydrate-active enzyme (CAZyme) families with known plant cell wall substrates (Cu=cutin). **b** Abundance matrix of candidate secreted effector proteins (CSEPs) that could be matched to experimentally verified genes in PHI-base. Genes are grouped based on knockout mutant phenotypes curated in PHI-base (Loss path=loss of pathogenicity). Both the CSEP orthogroup and name of the matched PHI-base gene are show on the x axis. ▼

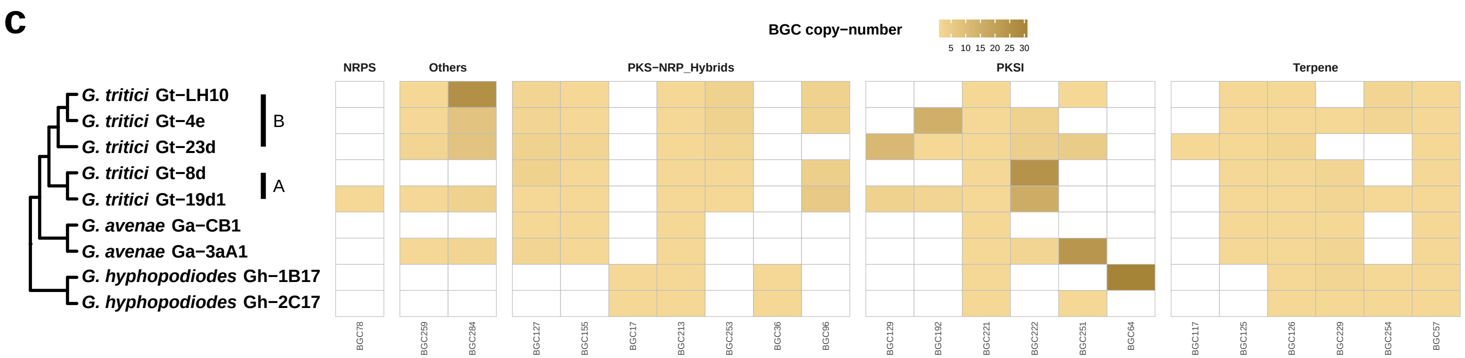


**Fig. S9 continued c** Abundance matrix of biosynthetic gene clusters (BGCs) belonging to classes assigned by BiG-SCAPE (NRPS= nonribosomal peptides synthetase, PKS=polyketide synthase, PKSI=type 1 polyketide synthase).

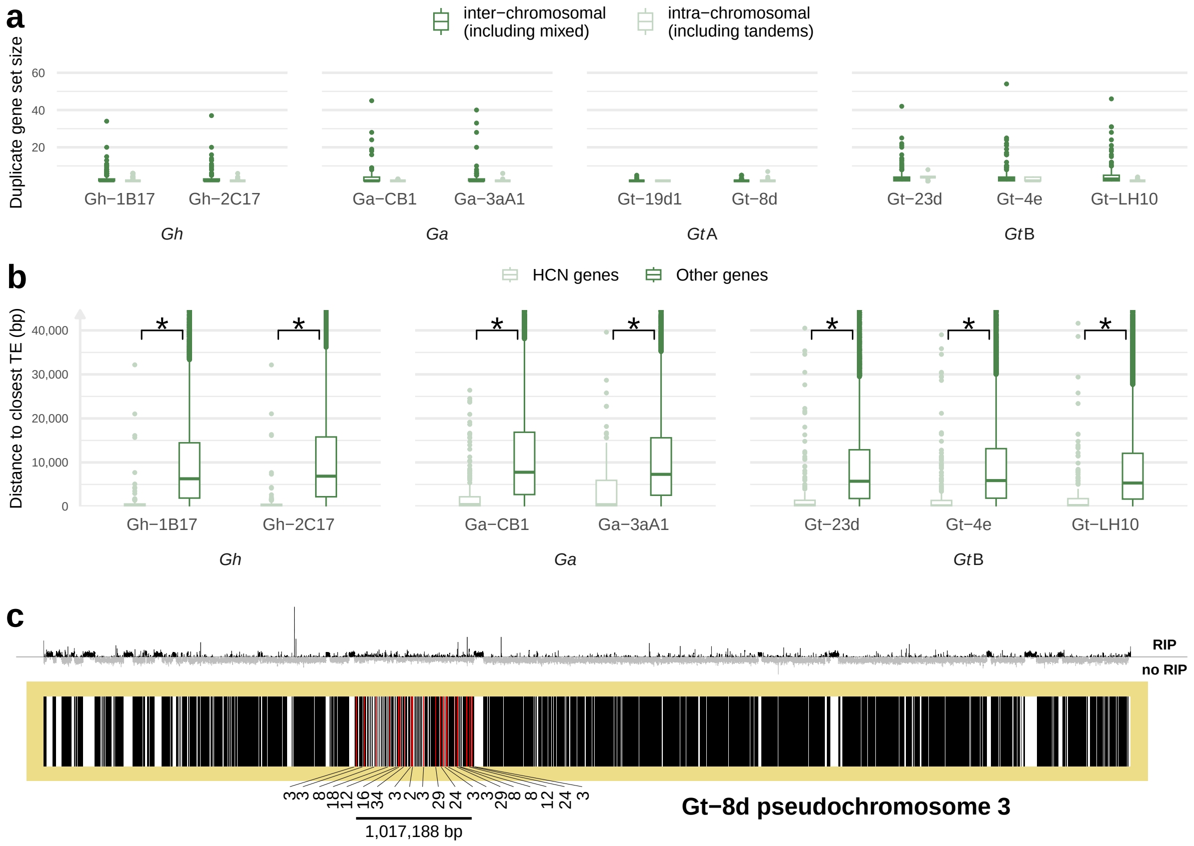


**Fig. S10 a** Duplicate gene set size for each strain categorised by whether the duplicates only occur on the same pseudochromosome (intra-chromosomal) or at least one duplicate is on a different pseudochromosome (inter-chromosomal). **b** Box plots showing the distance of high copy-number (HCN) genes to the closest transposable element (TE) versus other genes, with outliers > 40,000 bp distant not shown to improve visibility. An asterisk indicates where a Wilcoxon rank sum test found the mean TE distance to be significantly different for HCN genes versus other genes. **c** Region of *Gt* type A strain Gt-8d where low-copy orthologues of genes undergoing copy-number expansions in other lineages cluster on pseudochromosome 3. Red bars indicate the low-copy orthologues, with orthologue ID labelled below, and black bars indicate other genes. The composite RIP index (CRI) calculated in 500 bp windows is indicated above, coloured black when positive and thus indicative of RIP.

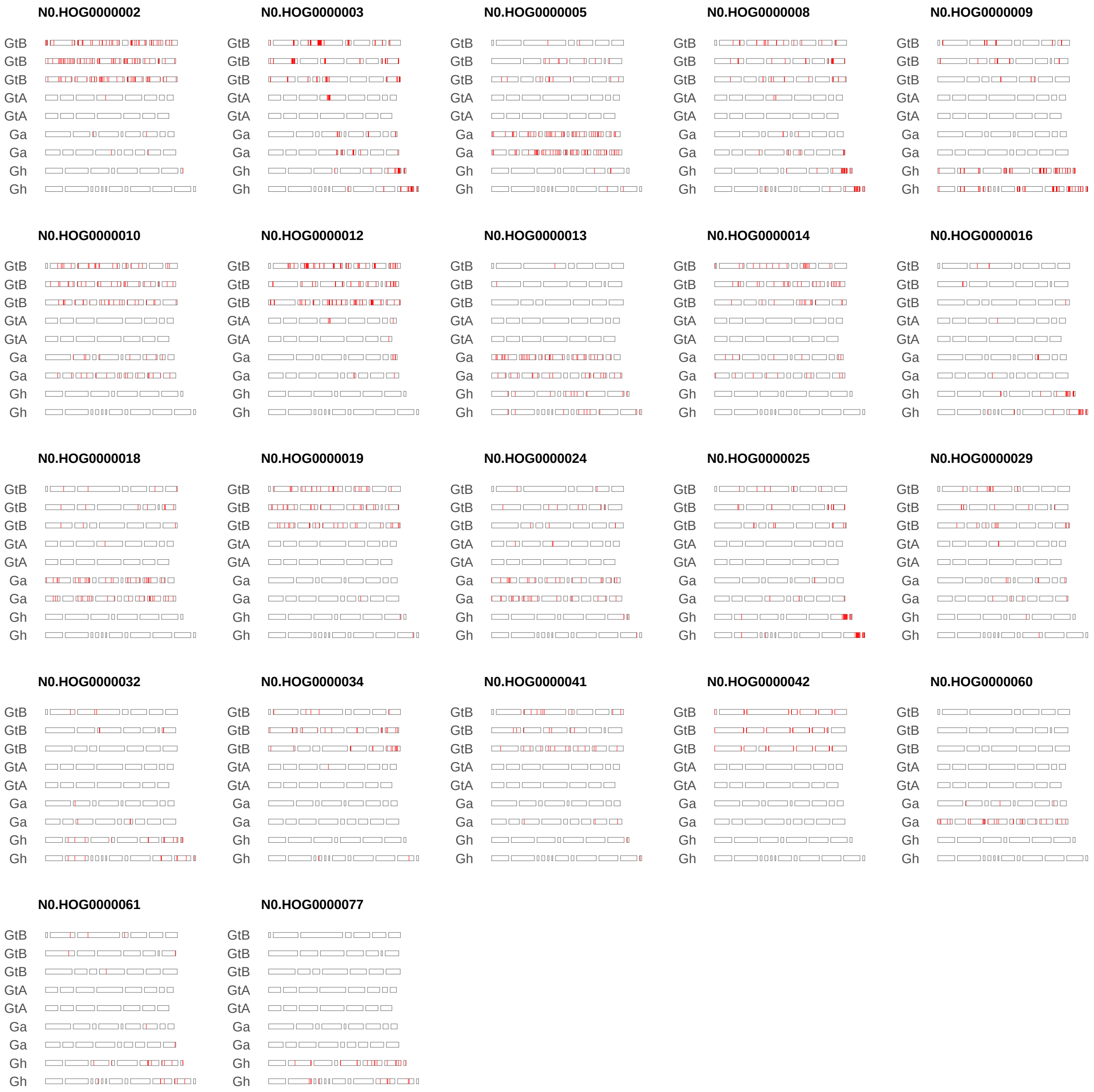


**Fig. S11** The location of copy-number expansions for 22 high-copy number genes. Red lines indicate the location of duplicates for the gene, and fragments are ordered syntenically according to GENESPACE (Fig. 2).

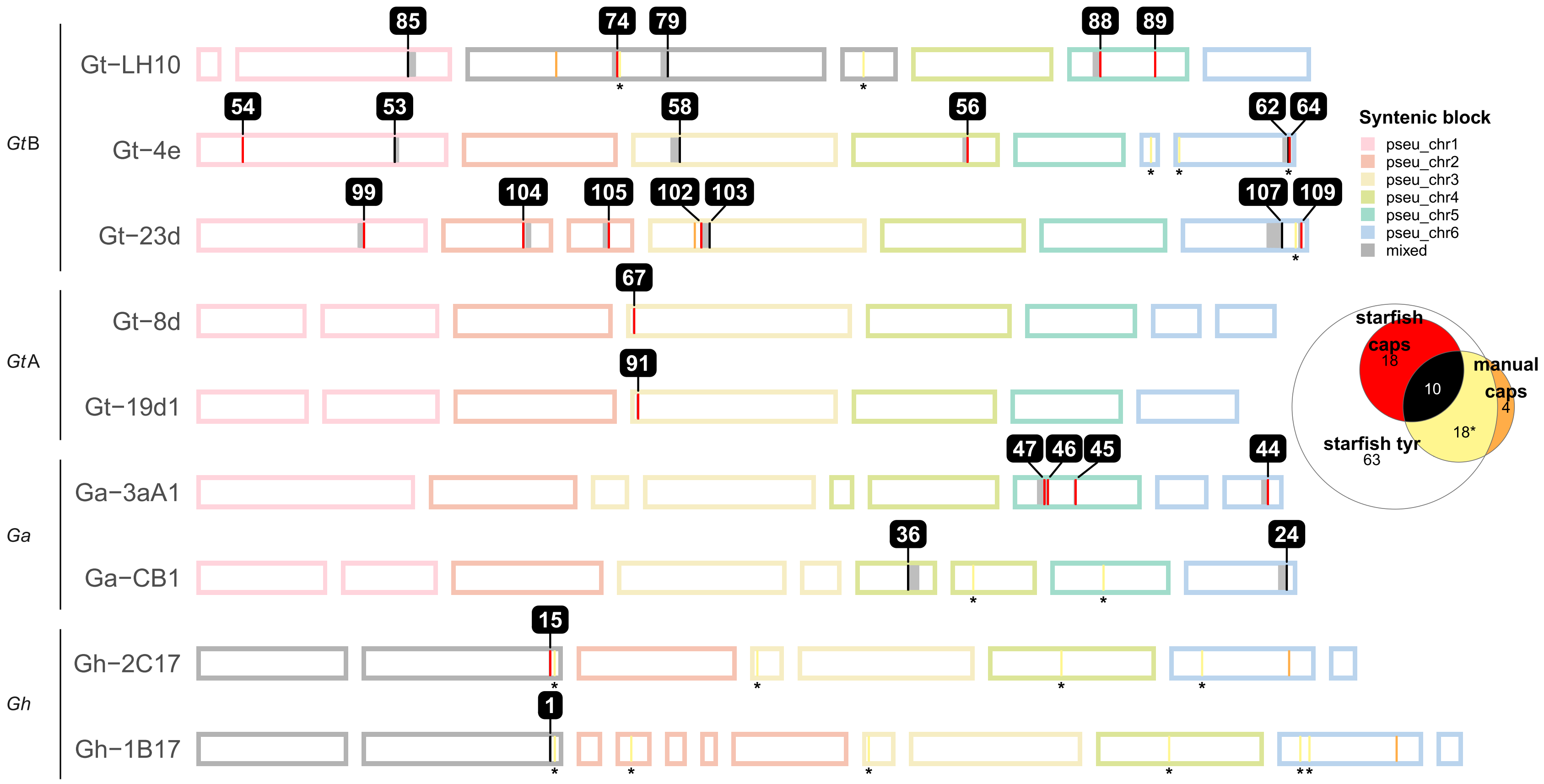


**Fig. S12** Location of *Starship* mobile element captain genes, with colour distinguishing whether genes were identified manually or using starfish (see inset euler plot). Grey blocks indicate associated cargo genes identified by starfish, and asterisks highlight manually identified genes which starfish categorised as non-captain tyrosine recombinases. Numbering corresponds to element IDs shown in Fig. 5b.

**a**

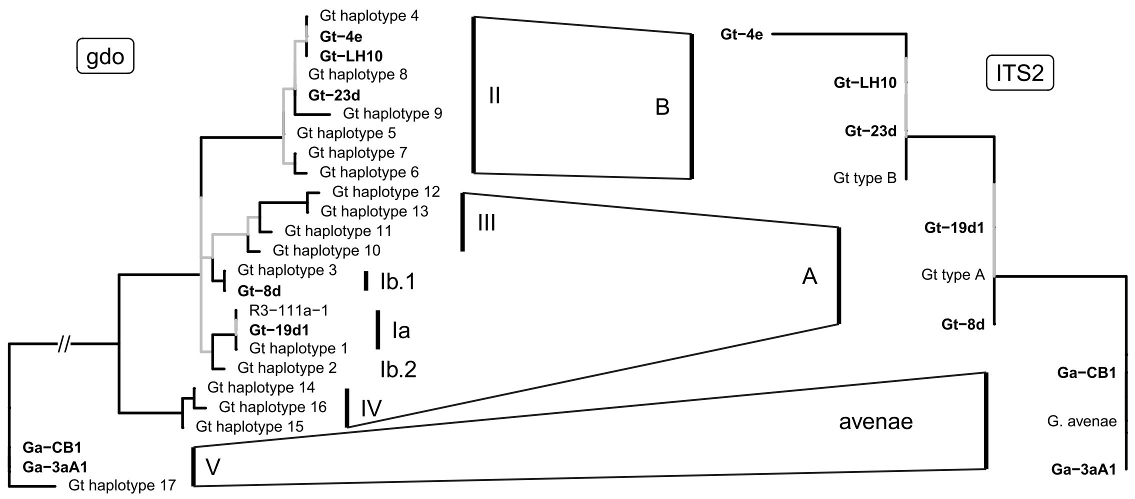


**b**

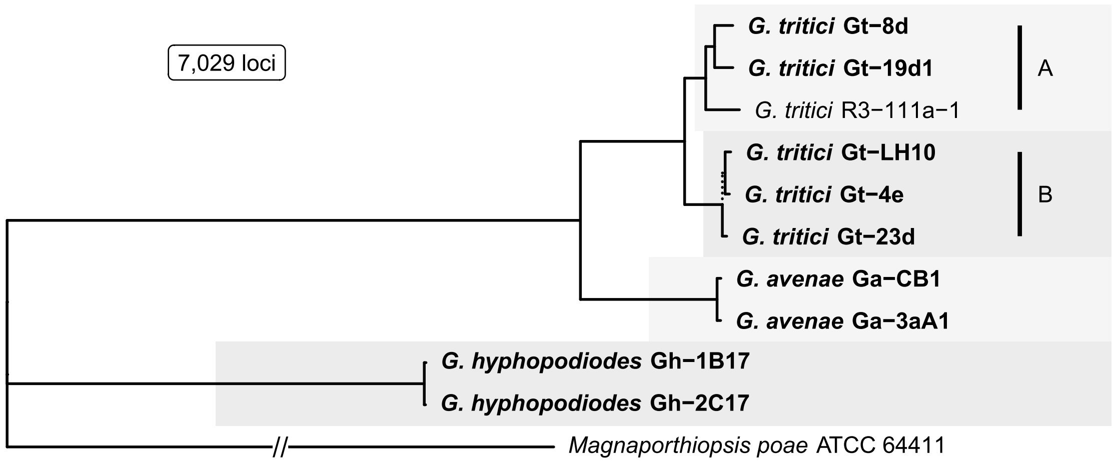


**Fig. S13 a** Gene trees for the gdo and ITS2 markers and their corresponding genetic groupings of *Gaeumannomyces tritici* (*Gt*) and *G. avenae*. Strains sequenced in this study are in bold and grey branches indicate <70 bootstrap support values. **b** *Gaeumannomyces* species tree with bars indicating type A and type B genetic groups for *Gt*. Strains sequenced in this study are in bold and dashed branches indicate <70 bootstrap support values.

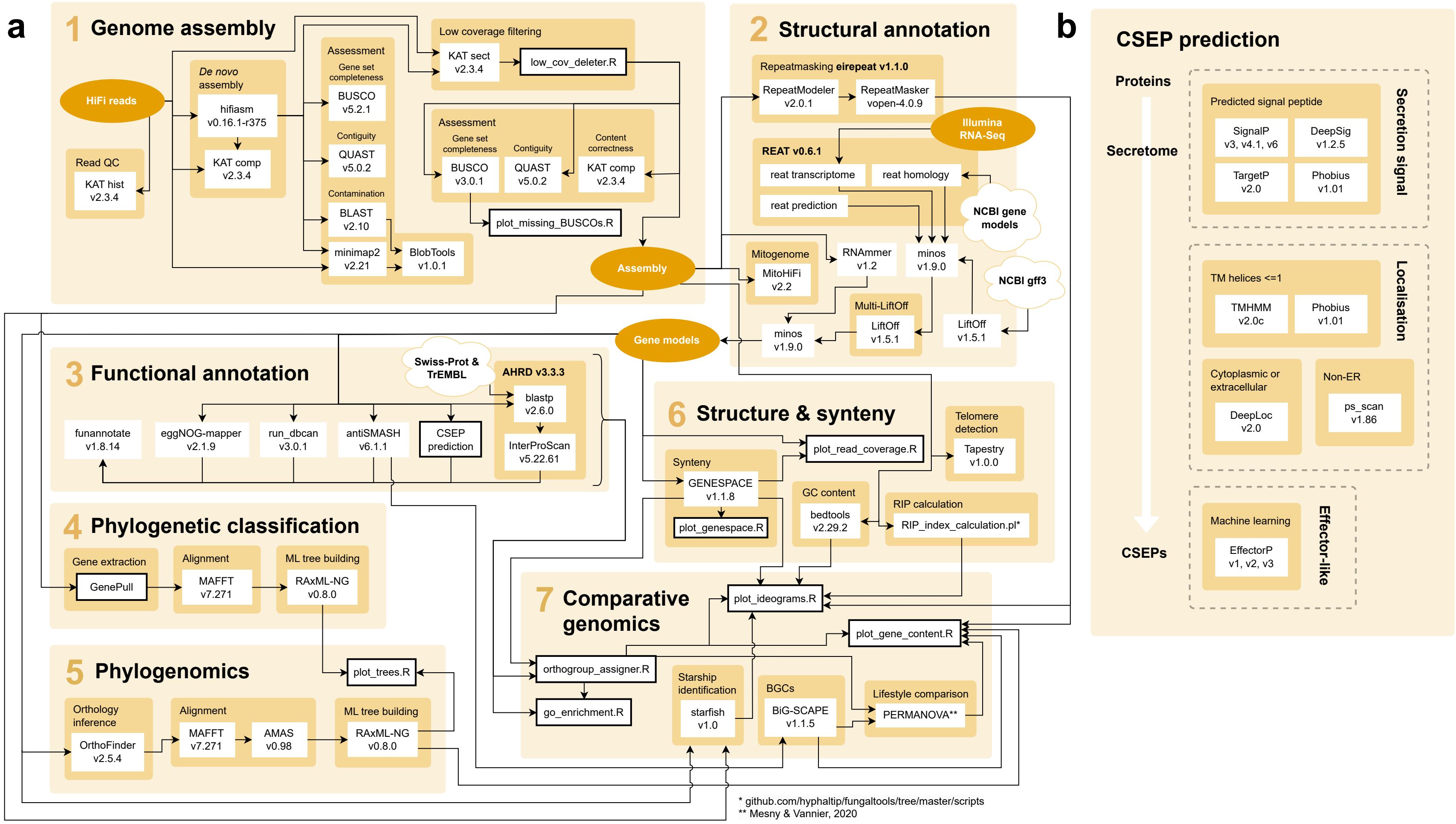


**Fig. S14** **a** Schematic summarising the bioinformatics analysis workflow developed in this study, available at https://github.com/Rowena-h/GaeumannomycesGenomics. Boxes outlined in black indicate custom scripts written for this study. **b** Summary of the steps involved in the candidate secreted effector protein (CSEP) prediction.

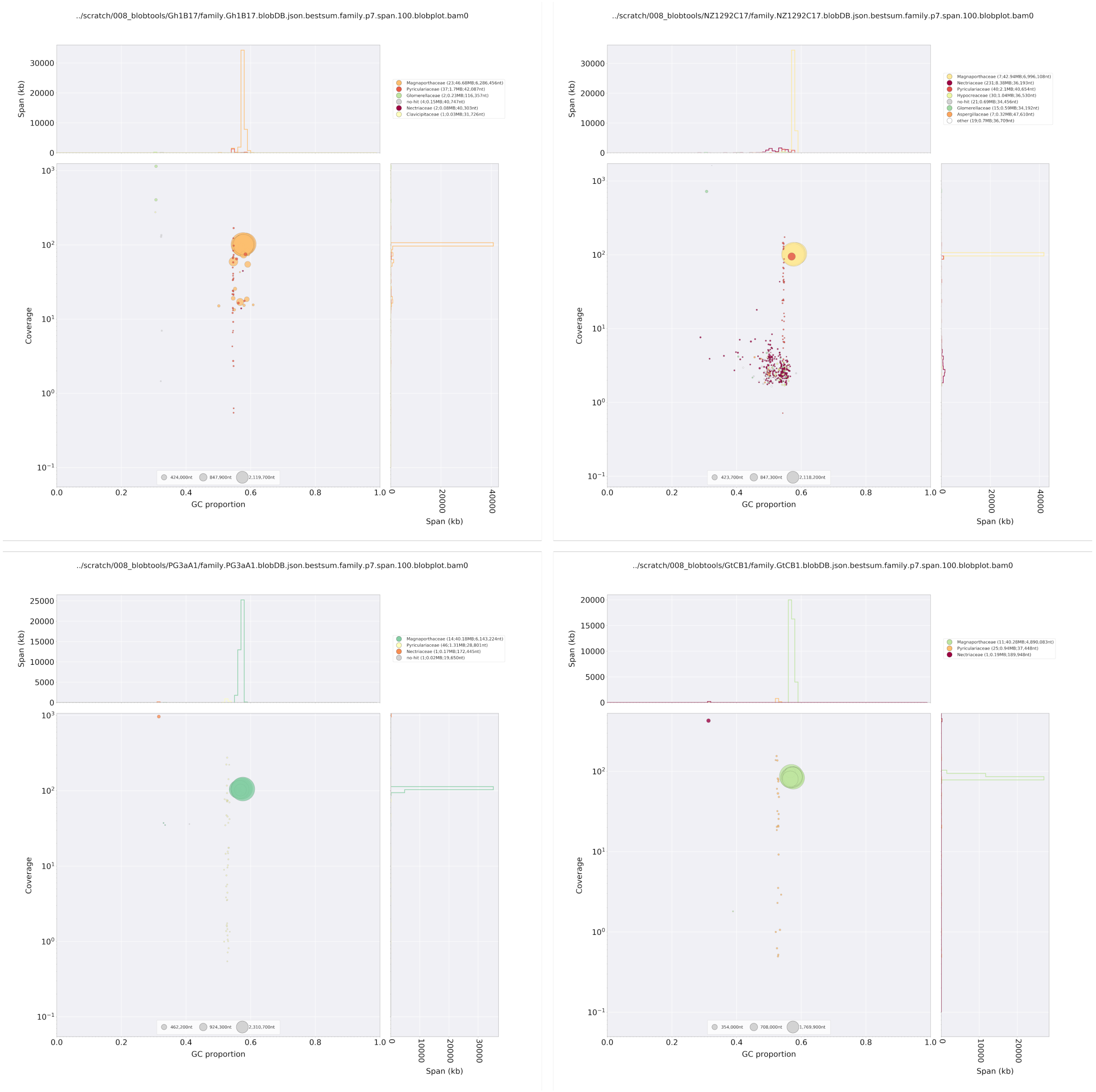


**Fig. S15** BlobPlots showing the taxonomic classification of reads based on coverage and GC content for each strain. From top left to bottom right: Gh-1B17, Gh-2C17, Ga-3aA1, Ga-CB1. ▼

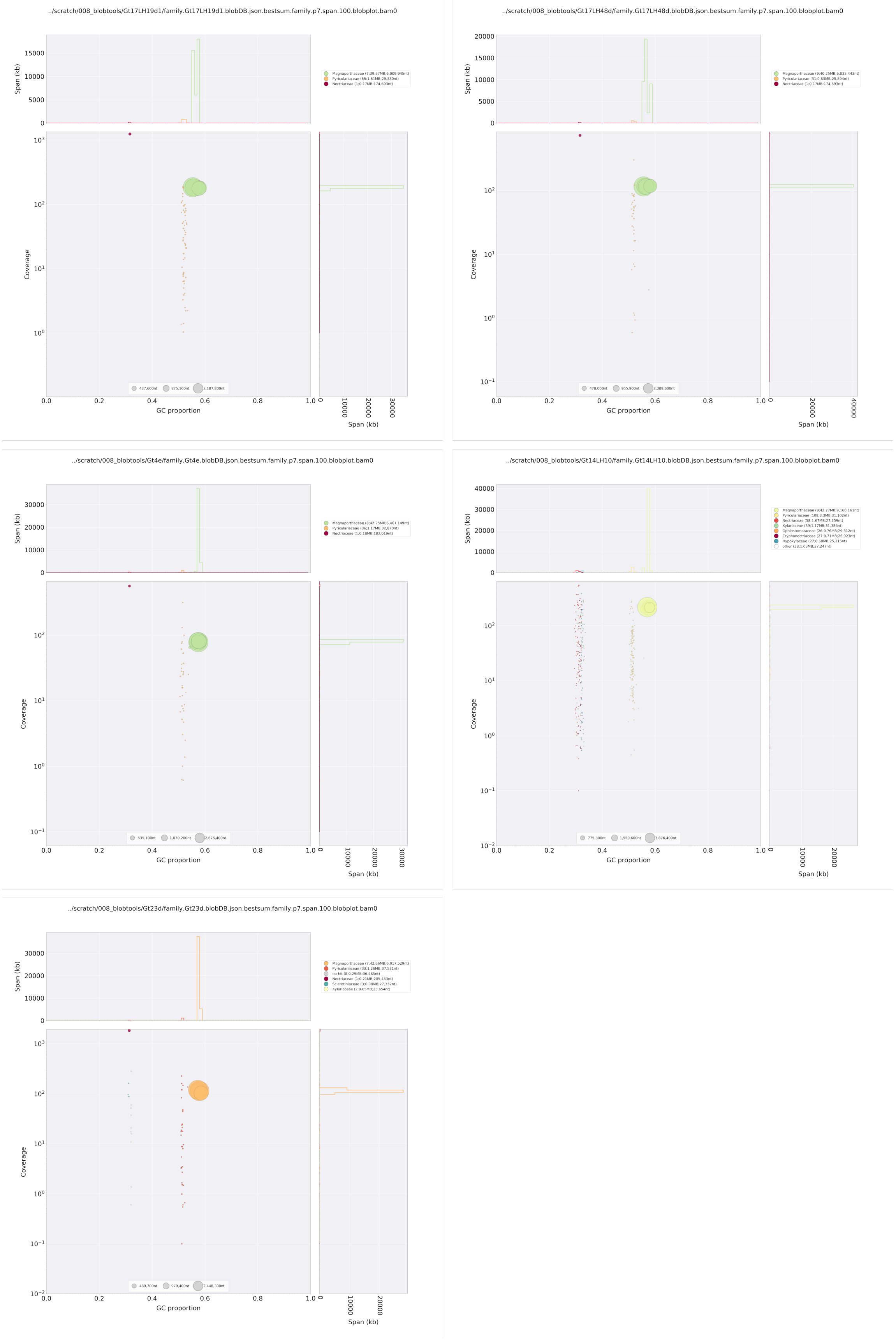


**Fig. S15 continued** From top left to bottom right: Gt-19d1, Gt-8d, Gt-4e, Gt-LH10, Gt-23d.

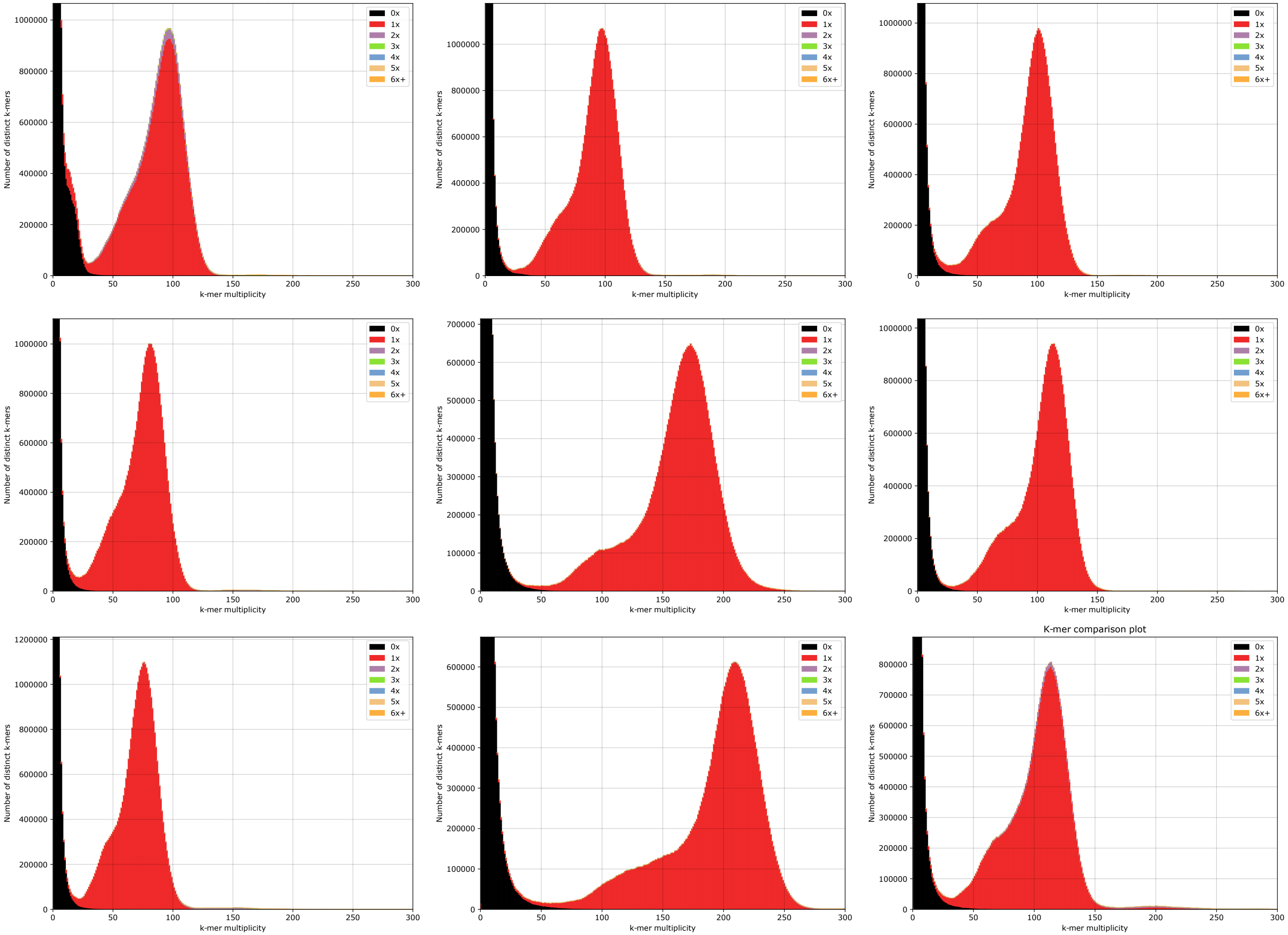


**Fig. S16** KAT COMP plots showing content correctness with respect to the input HiFi reads for each strain. From top left to bottom right: Gh-1B17, Gh-2C17, Ga-3aA1, Ga-CB1, Gt-19d1, Gt-8d, Gt-4e, Gt-LH10, Gt-23d.

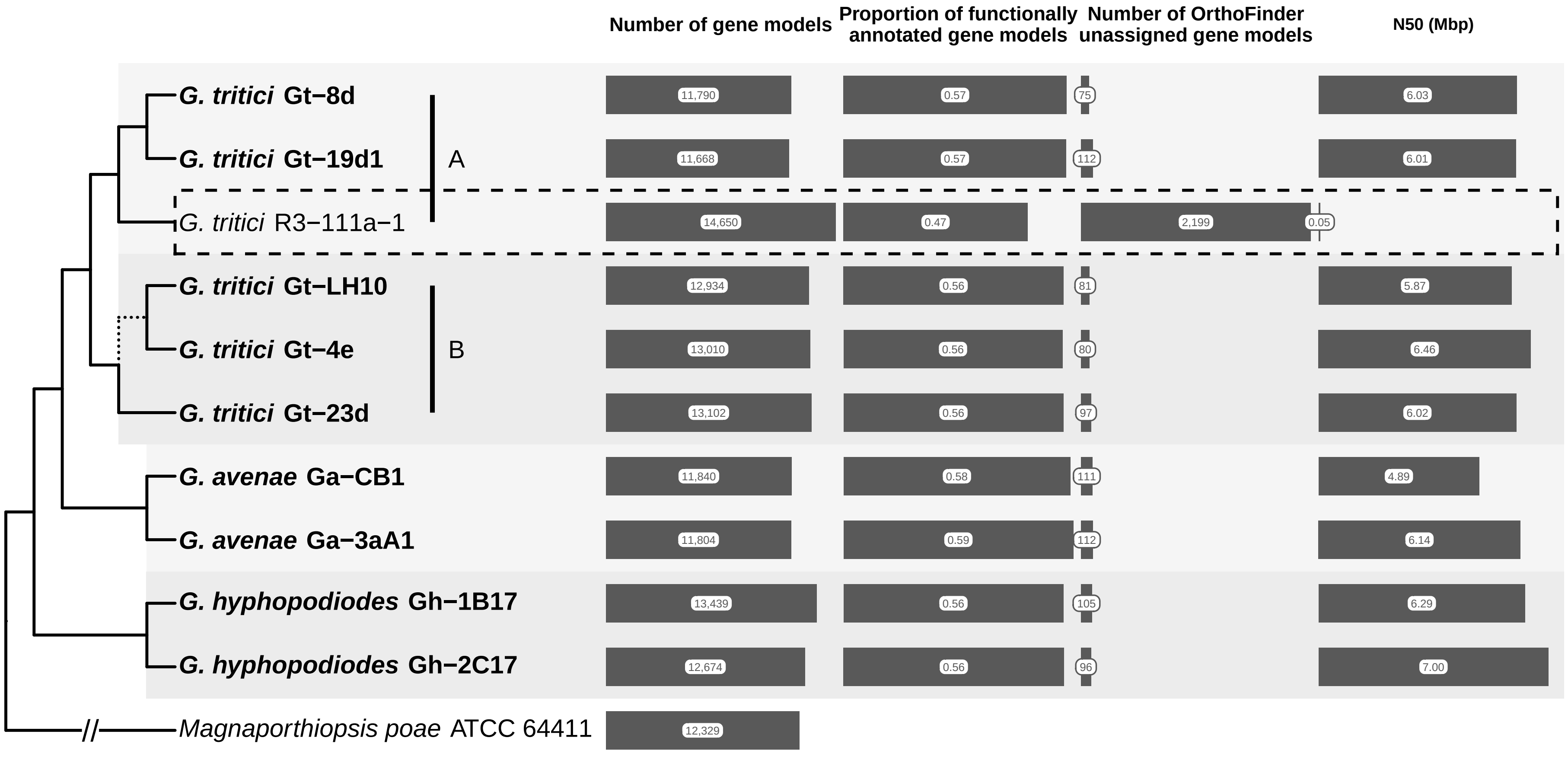


**Fig. S17** Comparison of the existing *Gaeumannomyces tritici* R3-111a-1 assembly/annotation (GCF_000145635.1) and the *Gaeumannomyces* assemblies/annotations generated in this study (bold). The proportion of functionally annotated genes refers to results from AHRD.
